# Predictable modulation of a spontaneous post-translational modification in living cells

**DOI:** 10.1101/2025.05.28.656619

**Authors:** Meghan S. Martin, Nomindari Bayaraa, Brittany T. Fox, Yu-Shan Lin, Rebecca A. Scheck

## Abstract

Glycation is a non-enzymatic post-translational modification associated with aging and disease. Because it occurs spontaneously, it is extremely difficult to control the extent of glycation at distinct sites within target proteins, especially in cellular systems. Here we report a chemical approach, referred to as ‘dialAGE’, that enables the site-specific control of protein glycation. This unique tool requires the introduction of just a single point mutation that modulates the glycation susceptibility of a nearby arginine. As proof-of-concept, extensive mass spectrometry analysis was performed to confirm that dialAGE can modulate site-specific glycation levels at multiple arginine residues in ubiquitin *in vitro*, enabling both enhanced and diminished glycation. Introduction of dialAGE point mutations and/or glycation with the biologically relevant glycating agent methylglyoxal did not affect polyubiquitin chain formation using *in vitro* ubiquitination assays. Furthermore, we show that dialAGE can be used to modulate ubiquitin glycation levels in living mammalian cells. We therefore anticipate that this method will be particularly useful for enabling the study of glycation as a genuine, functional, post-translational modification.

## Introduction

Sugars and sugar-derived metabolites—aldehydes and ketones—react spontaneously with amino and guanidino groups on proteins.^[1–3]^ The resulting post-translational modification (PTM) is known as glycation (or ‘non-enzymatic glycosylation’), which produces a remarkably large, diverse set of adducts collectively known as advanced glycation end-products (AGEs).^[4]^ Though protein glycation has been correlated with aging and disease ever since its discovery,^[1–3]^ it still remains unclear whether glycation is only a marker for metabolic dysregulation or could also have a causal role in disease etiology.^[4,5]^ Without ‘writer’ or robust ‘eraser’ enzymes to manipulate, this question persists because the glycation of specific proteins has been nearly impossible to control. The vast majority of cellular and *in vivo* experiments broadly induce glycation by increasing the concentrations of reactive metabolites through treatment with high levels of glucose (or other glycating agents), inhibition of glyoxalase enzymes that detoxify the potent glycating agent methylglyoxal (MGO), or induction of hyperglycemia through high glycemic index diets.^[4–10]^ Each of these approaches causes widespread changes in both signaling and glycation, making it extraordinarily difficult to attribute specific glycation events to a given outcome.

Although glycation occurs spontaneously, it is also known to transpire selectively.^[11–14]^ Despite numerous potential reactive sites, namely Arg and Lys side chains and the N-terminus, only a subset are glycated.^[6]^ Past work has suggested that this observed selectivity is due to increased solvent accessible surface area^[11,14]^ or decreased Arg/Lys pK*_a_*.^[14–17]^ However, recent work,^[18]^ including our own,^[12,13]^ has demonstrated that the proximal residues surrounding glycation sites are major drivers for glycation. In particular, we have shown that nearby tyrosine and polar residues enhance glycation on short, unstructured peptides, while surrounding clusters of acidic residues inhibit it. ^[12,13]^

Our inspiration to precisely control site-specific glycation events stems from the longstanding observation that the surrounding microenvironment is known to modulate pK*_a_* and/or nucleophilicity for a given site within a structured protein. While this phenomenon is most appreciated within enzyme active sites,^[19]^ it has also been exploited for activity- and reactivity-based protein profiling^[20–23]^ and regioselective protein bioconjugation.^[24–42]^ Indeed, there are many examples of regioselective labeling of amino groups, like Lys side chains and protein N-termini.^[25–32]^ Though chemoselective modification of the Arg guanidino group is less common, it has also been accomplished in a site-selective fashion.^[33–35]^ However, there are only rare instances where a specific chemical environment has been tuned, designed, or engineered to enable site-selective chemical labeling of proteins.^[36–42]^ Additionally, to our knowledge, such an approach has so far been mostly limited to synthetic protein labeling methods and has not been applied to modulate levels of naturally occurring PTMs.

In this work, we thoughtfully incorporate the impactful sequence elements that we have shown to influence glycation for peptides within a full-length, folded protein, enabling the modulation of glycation at specific sites. This strategy, which we call “dialAGE”, is uniquely suited to tune AGE levels at specific sites on specific proteins. Using mass spectrometry, we demonstrate that dialAGE can be used to selectively tune the reactivity of multiple discrete glycation sites within the small protein ubiquitin (Ub). Our results show that dialAGE does not affect polyubiquitin chain formation and can be applied both *in vitro* and in living mammalian cells. Our findings suggest that dialAGE is a powerful new tool for controlling glycation that will greatly advance the study of glycation biology.

## Results and Discussion

To evaluate the feasibility of dialAGE, we selected the small protein ubiquitin (Ub) as an initial target due to its stability, well-understood structure, and substantial biological and disease relevance. Several reports have suggested that glycation adversely affects the ubiquitin proteasome system (UPS), though the exact mechanism is not understood.^[43–47]^ Additionally, our previous work showed that Ub was among the proteins most extensively glycated by MGO *in vitro*,^[12]^ leaving open the possibility that it could be a target for glycation *in vivo*. Consistent with prior reports, we found that MGO—a reactive α-oxoaldehyde—exhibited chemoselectivity for Arg,^[12]^ and produced multiple AGEs with known structures (Figure 1A). Although the majority of Ub biology focuses on its seven Lys residues, it also has four Arg residues (R42, R54, R72, and R74), that can influence polyubiquitin chain formation, disassembly, and quality control.^[48–51]^ In particular, R42 is reported to interact with Ub conjugating enzymes,^[51,52]^ other Ub-dependent recognition enzymes,^[53]^ and is the site of phosphoribosyl-linked ubiquitination that has been shown to inhibit the UPS during *L. pneumophila* infection.^[54]^ Therefore, we first chose to prepare dialAGE^R42^ Ub variants that we predicted to exhibit varying susceptibility to glycation at R42.

**Figure 1.**
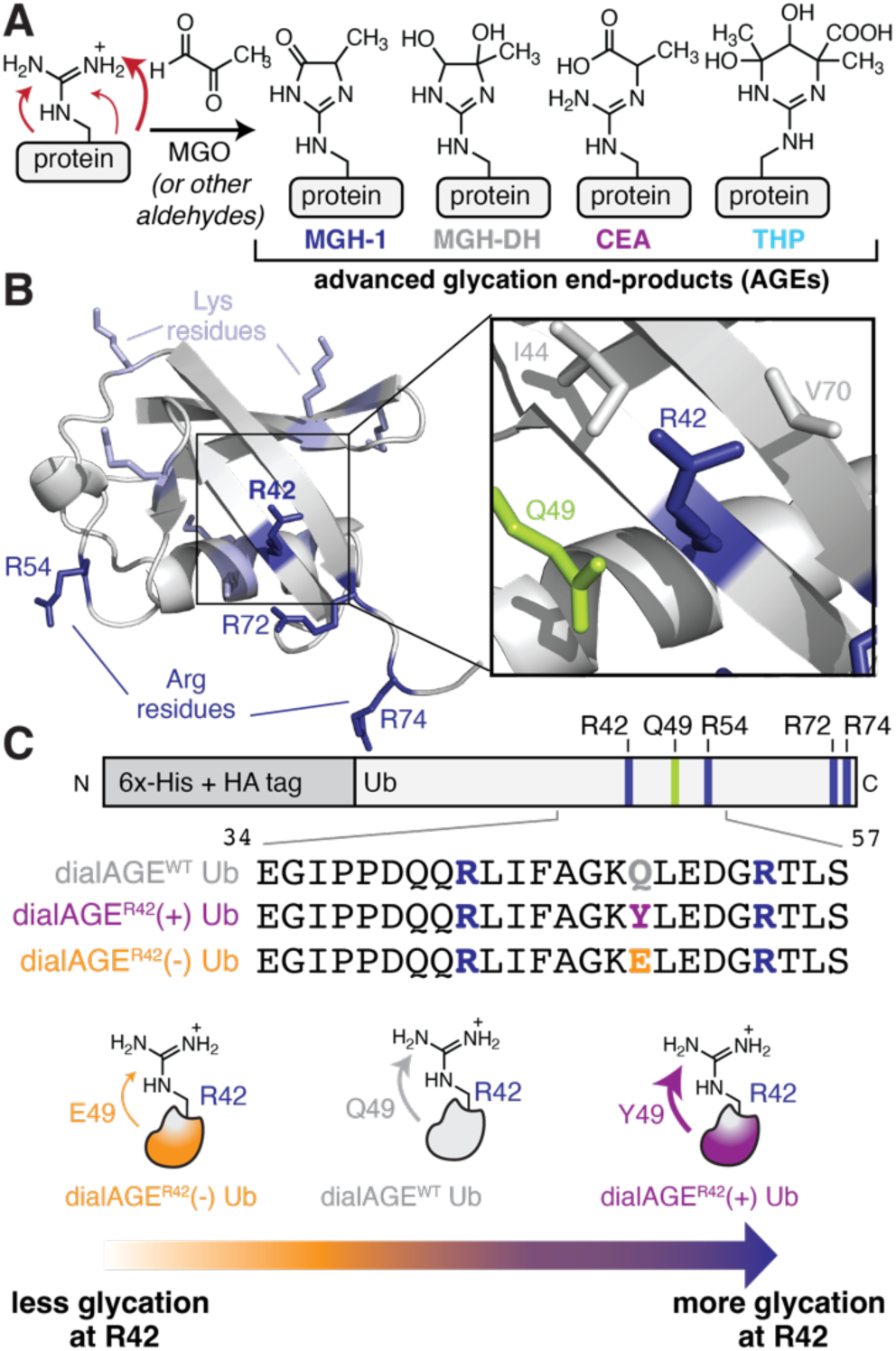
Designing a strategy to modulate site-specific glycation of ubiquitin (Ub). A) Glycation is a non-enzymatic PTM in which aldehyde-containing metabolites, such as methylglyoxal (MGO), react with Arg or Lys side chains to produce a set of modifications known as advanced glycation end-products (AGEs). MGO-derived AGEs include hydroimidazalone isomers such as MGH-1, the dihydroxyimidazolidine (MGH-DH), carboxyethyl-arginine (CEA), and tetrahydropyrimidine (THP). B) To identify sites close enough in space to modulate glycation at R42, the crystal structure of ubiquitin (1UBQ) was closely inspected, showing the side chains of I44, Q49, and V70 to be within 5Å of the guanidinium sidechain of R42. C) To evaluate the impact of Q49 substitutions on R42 glycation, “dialAGE^R42^” variants were made by mutating Q49 to Tyr for a dialAGE^R42^(+) variant and to Glu for a dialAGE^R42^(−) variant. The dialAGE^WT^ variant includes the wild-type Ub sequence fused to N-terminal 6×-His and HA epitope tags.

To develop dialAGE^R42^, our first goal was to identify the surrounding residues close enough in space to influence R42 glycation. Careful inspection of the Ub crystal structure (Figure 1B) revealed I44, Q49, and V70 to be within 5Å of the guanidinium sidechain of R42. As I44 and V70 are both part of the critical Ub hydrophobic core, we opted to mutate Q49, located in a nearby β-strand. We prepared dialAGE^R42^(+) (Q49Y) and dialAGE^R42^(−) (Q49E) Ub variants, which we expected to increase or decrease glycation compared to dialAGE^WT^ (wild-type), respectively (Figure 1C). Because protein structure can influence glycation,^[12]^ we used circular dichroism (CD) spectroscopy to confirm that all dialAGE Ub variants exhibited highly similar folds (Supporting Information, Figure S1). This result suggests that any potential observed differences in glycation could be attributed to the change in local environment introduced by the dialAGE^R42^ mutations, rather than by a change of the folded structure.

To assess glycation, 50 *μ*M of each dialAGE Ub was incubated with 100 *μ*M MGO at 37 °C (Figure 2A,B; Supplementary Information, Figures S2, S3, S4). After 24 h, total glycation was assessed using intact protein LC-MS. We estimated differences in glycation by quantifying the remaining unmodified Ub after treatment. This revealed statistically significant differences for dialAGE^WT^ (42.5% unmodified), dialAGE^R42^(+) (30.7% unmodified), and dialAGE^R42^(−) (56.3% unmodified) Ub (Figure 2B). These differences persisted after 48 h of MGO treatment, with more glycation observed for dialAGE^R42^(+) (27.0% unmodified) than for dialAGE^WT^ (39.8% unmodified), and dialAGE^R42^(−) (48.5%) (Supporting Information, Figure S4). These results suggest that the dialAGE strategy modulates total glycation levels for protein targets *in vitro*.

**Figure 2.**
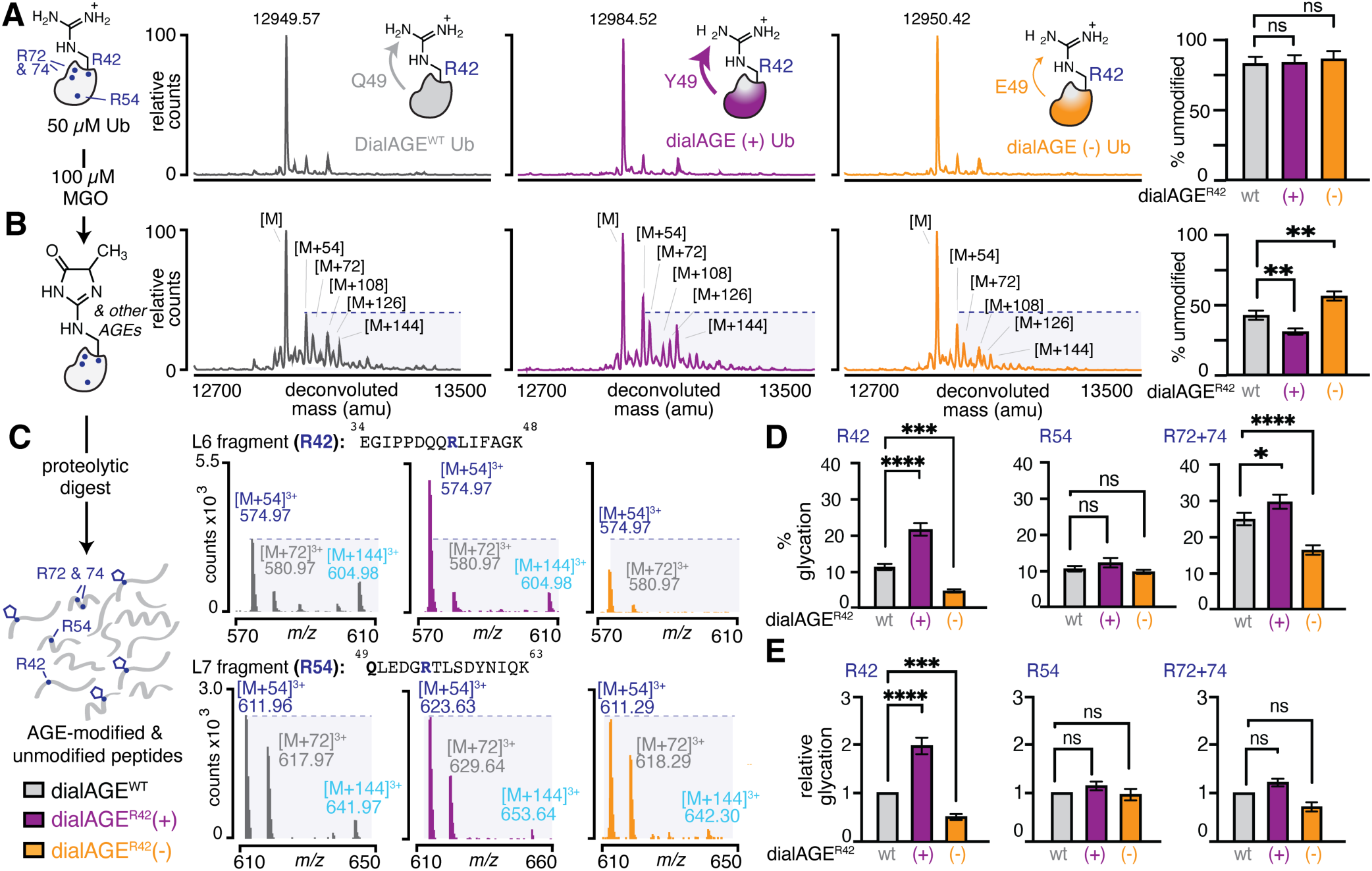
Revealing site-specific control of glycation with dialAGE. A,B) To evaluate Ub glycation, dialAGE^WT^ and dialAGE^R42^ variants (50 *μ*M) were treated with MGO (100 *μ*M; 2 equiv.) for 24 h at 37 °C in phosphate buffered saline at pH 7.4, leading to statistically significant differences in total glycation as observed by intact mass spectrometry. The extent of modification was determined by quantifying the intensity of the remaining unmodified peak ([M]) (A) before and (B) after MGO treatment. C) Following digestion with Lys-C, it was possible to observe dialAGE-dependent differences in glycation for R42 (L6 fragment) by comparing peak intensities in the resulting mass spectra. These differences were not present for R54 (L7 fragment). Total D) absolute or E) relative glycation levels were quantified for each dialAGE variant at each Arg. This quantification shows significantly modulated glycation at R42, and to a lesser extent R72+74, but constant levels of glycation at R54. Bar graphs represent mean±SEM. Nondirectional (two-tailed) two way ANOVA using Tukey’s multiple comparison tests were used to determine if each variant yielded statistically significant differences in glycation. P<.05 (*), p<.01 (**), p<.001 (***), p<.0001 (****).

Our next goal was to determine if these observed differences in glycation were site-specific or due to a modulation of reactivity across multiple sites. To assess this, we used proteolytic digest by Lys-C (Figure 2C-E, Supporting Information, Figures S5 & S6) or trypsin (Supporting Information, Figure S7) followed by LC-MS/MS analysis comparing relative glycation at each Arg after MGO treatment. For R42, this revealed that dialAGE^R42^(+) exhibited an approximately 2-fold increase while dialAGE^R42^(−) Ub exhibited a roughly 2-fold decrease in glycation relative to dialAGE^WT^ (Figure 2C-E). Levels of glycation at R54 remained virtually identical for dialAGE^R42^(+), dialAGE^R42^(−) and dialAGE^WT^ (Figure 2C-E). Similar results were observed when using trypsin instead of Lys-C (Supporting Information, Figure S7). Time course studies revealed that even at the earliest timepoint (1 h), there were dialAGE-dependent differences in glycation levels at R42, which grew more pronounced at later time points (Supplementary Information, Figures S8 & S9). Moreover, there were no apparent differences in glycation at R54 at any time point. The remaining two Arg, R72 and R74, appear in the same Lys-C fragment and could not be reliably disambiguated by MS/MS. Analysis of their glycation was grouped, revealing only a minor increase (1.2 fold) in glycation for dialAGE^R42^(+) and a minimal decrease (0.7 fold) for dialAGE^R42^(−) as compared to dialAGE^WT^ (Figure 2D,E; Supporting Information, Figure S6,S9). We suspect that these differences in glycation, though smaller than the ones observed at R42, are likely due to the location of Q49, which is 5 Å from R42 but is still somewhat proximal (6.3 Å) to R72. As expected, we note that we did not observe any glycation for the seven Lys residues (or Ub N-terminus) under these conditions.

Glycation occurs through a multi-step process and typically produces multiple AGEs even on a single glycation site.^[4]^ We have previously shown that discrete AGEs can interconvert,^[13]^ suggesting that dialAGE could modulate not only the total extent of glycation observed at a single site, but also the specific distribution of AGEs that form. To assess this, we quantified AGE distributions for each dialAGE Ub variant after 24 h (Supporting Information, Figure S10). No major differences were observed in the proportional AGE distributions between variants at R42. An [M+54] adduct dominated, comprising 73.0 ± 1.1%, 73.9 ± 5.4%, and 84.2 ± 4.8% of all AGEs for dialAGE^WT^, dialAGE^R42^(+), and dialAGE^R42^(−) respectively. The next most prevalent was an [M+72] adduct, which accounted for 18.9 ± 1.2%, 19.2 ± 4.3%, and 12.7 ± 3.9% of AGEs, followed by small amounts (8.1 ± 1.5%, 6.8 ± 3.0%, and 3.1 ± 1.3%) of an [M+144] species. We also found that the AGE distribution at R54 was nearly identical to that observed for R42 and was consistent across all dialAGE variants. Distributions of R72+R74 adducts also appeared nearly identical between variants. Similarly, at early timepoints (1, 3, and 5 h) and 48 h we also observed no major differences in AGE distributions across all Arg (Supporting Information, Figure S10). These results suggest that dialAGE influences the rate of AGE formation for a given site.

To evaluate the scope of dialAGE, we sought to incorporate additional point mutations that we expected to modulate R42 in a predictable way. In our prior work, we found that the features that increased glycation levels included the presence of Tyr and polar residues, like Ser, though Tyr appeared to contribute more favorably to glycation than any other residues.^[12,13]^ Additionally, negatively charged amino acids like Asp could elicit a similar, if not greater, decrease in glycation compared to Glu.^[12,13]^ Therefore, we prepared an additional set of dialAGE^R42^ Ub variants: dialAGE^R42^(+)^Q49S^ and dialAGE^R42^(−)^Q49D^ to increase or decrease R42 glycation, respectively. Following *in vitro* glycation, we observed a clear decrease in total protein glycation for dialAGE^R42^(−)^Q49D^ compared to dialAGE^WT^, while there were no obvious differences for dialAGE^R42^(+)^Q49S^. On a site-specific level, dialAGE^R42^(−)^Q49D^ resulted in a roughly 3-fold decrease in R42 glycation, which was even greater than the extent of decrease we observed with our original dialAGE^R42^(−)^Q49E^ variant. Similarly, we observed consistent levels of glycation at R54 and R72+74 compared to dialAGE^WT^. For the dialAGE^R42^(+)^Q49S^ variant, we observed a small, but statistically insignificant, increase in glycation at R42, and there were no changes in glycation levels at any other Arg (Supporting Information, Figure S11). Both variants also produced a highly similar distribution of AGEs to dialAGE^WT^, though we note that irrespective of the observation that dialAGE^R42^(−)^Q49D^ did not form any [M+144], the least abundant AGE observed for other dialAGEs. Taken together, these results confirm that the behavior of dialAGE matches closely with our previous work that has allowed us to learn the sequence features governing selective glycation events.

Our next goal was to develop a better understanding for how dialAGE affects the chemical environment surrounding R42, thereby influencing glycation. To do so, we performed molecular dynamics simulations using the Amber ff19sb forcefield, which has been benchmarked on several systems, including 1UBQ.^[55]^ For each variant, we performed five parallel sets of molecular dynamics simulations using the same initial structure, but different initial velocities (see Supporting Information). We evaluated several features as a function of the dialAGE mutation, including root mean square deviation (RMSD), root mean square fluctuation (RMSF), and radius of gyration, none of which showed any significant variation compared to wild-type Ub that could explain the observed differences in glycation (Supporting Information, Figure S12). We also found that there were no differences in the solvent accessible surface area (SASA) for R42, further corroborating our prior results.^[12]^

Encouragingly, the molecular dynamics simulations also provided insight into how dialAGE works, revealing that the distance between the terminal carbon of the R42 side chain and the terminal carbon of the side chain at site 49 was modulated by dialAGE^R42^ mutations (Figure 3A,B). For dialAGE^R42^(−)^Q49E^, there was a significant population shift towards shorter R42…49 distances that were less than 5Å, while for dialAGE^R42^(+)^Q49Y^, there was a noticeable shift towards longer distances that were greater than 5Å (Figure 3A, left). To compensate, other nearby residues (e.g., E51 and D52) that could interact with R42 also moved towards (dialAGE^R42^(+)) or away dialAGE^R42^(−)) from it (Figure 3A, middle and right; Figure S13). We also found introduction of dialAGE^R42^ mutations did not perturb the observed distances between R54 and its nearby residues, which is consistent with the site-specific experimental results we observed (Figure 2, Supporting Information, Figure S13). Representative structures depicting each of these populations show that the Q49E mutation likely works by trapping R42 in an electrostatic interaction that impedes glycation (Figure 3C). Conversely, Q49Y lengthens the distance between R42 and site 49. Based on our prior work, we expect that this not only provides room for MGO to access R42, but also likely positions the Tyr phenol in a location that enables it to facilitate glycation.

**Figure 3.**
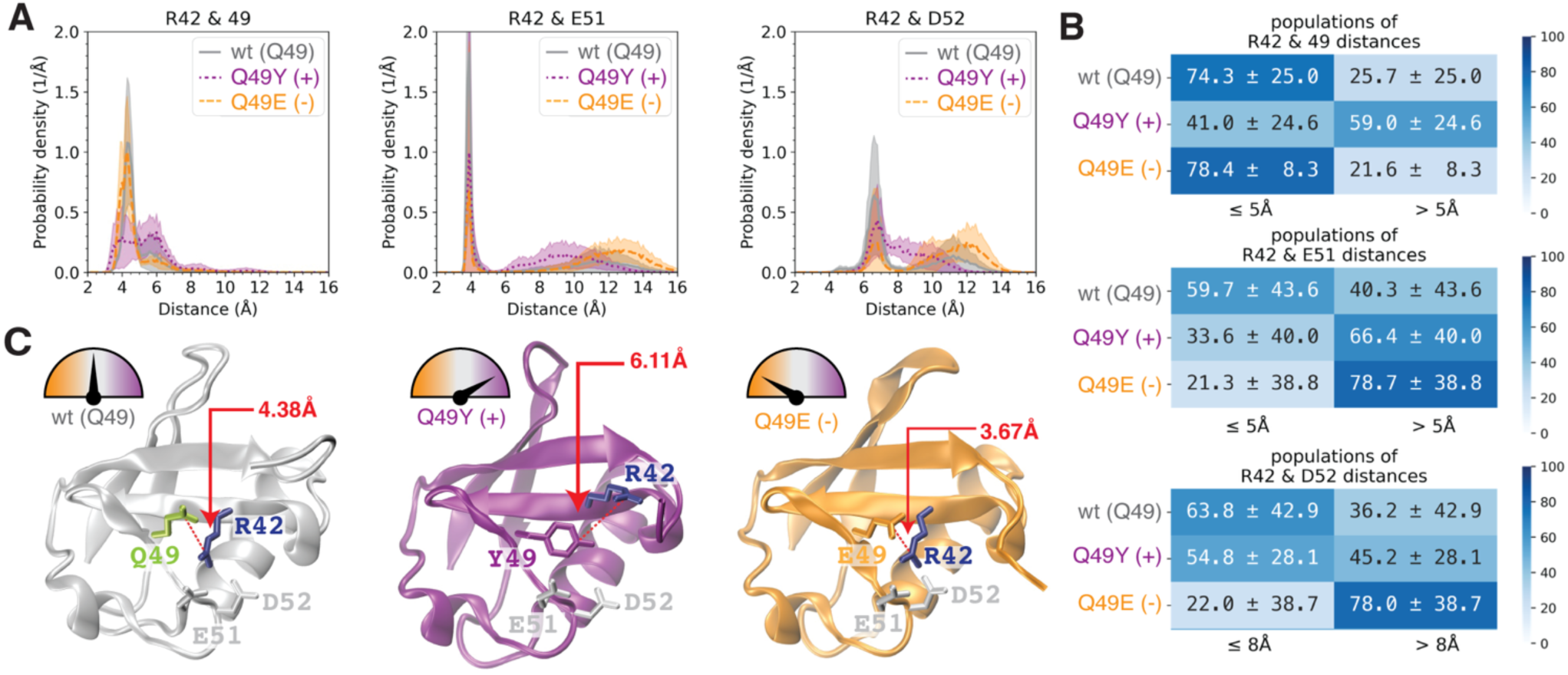
Evaluating dialAGE using molecular dynamics simulations. A) Distance distributions (with standard deviations in shaded regions) between the terminal carbon of the R42 side chain and the terminal carbon of the side chain at site 49 (left), between the terminal carbon of the R42 side chain and the terminal carbon of the E51 side chain (middle), and between the terminal carbon of the R42 side chain and the terminal carbon of the D52 side chain (right). B) Populations of conformations where the distances between the terminal carbon of the R42 side chain and the terminal carbon of a nearby side chain are above or below a given threshold: (left) ≤5Å for R42…X49 and R42…E51, ≤8Å for R42…E51; (right) >5Å for R42…X49 and R42…E51, >8Å for R42…E51. C) Representative structures illustrating the major population for wild-type, Q49Y (dialAGE^R42^(+)), and Q49E (dialAGE^R42^(−) Ub variants.

Having confirmed that dialAGE is a feasible strategy for influencing glycation at R42, we next sought to control glycation at R54, which sits in an unstructured loop in the Ub crystal structure. We therefore prepared dialAGE^R54^(+) and (−) variants by mutating T55 (Supporting Information, Figure S14), which is situated proximally to R54 in the available crystal structure (6.5Å). Following the same glycation and digestion protocols, we observed the expected increases and decreases in total glycation, as compared to dialAGE^WT^. Indeed, dialAGE^R54^(−) led to a statistically significant site-specific decrease in glycation at R54, and dialAGE^R54^(+) produced a statistically significant increase at R54. However, while the effect was site-specific for dialAGE^R54^(−), dialAGE^R54^(+) also produced a statistically significant increase in glycation for all Arg. In both cases, the AGE distributions at each Arg were highly similar to that observed for dialAGE^WT^ (Supporting Information, Figures S15 & S16). We also confirmed that both dialAGE^R54^ variants have nearly identical folds to dialAGE^WT^ at both 37 °C and 60 °C, suggesting that this behavior is not due to major differences in thermal stability (Supporting Information, Figure S17). To see if it was possible to similarly generate a global decrease in glycation, we prepared a dialAGE^R54^(−)^T55F^ variant, as our previous work showed that Phe could decrease glycation. We reasoned that the structural similarity between Phe and Tyr might produce global glycation modulation across all Arg, but with an opposite effect. However, we found no differences in glycation at any Arg with this variant (Supporting Information, Figure S18).

Because of the location of T55 within a turn-rich region,^[56]^ we suspected that this site may be less well-suited to dialAGE. Using molecular dynamics simulation, we found that the distances between the terminal carbon of the R54 side chain and the terminal carbon of the side chain at site 55 were substantially larger (9Å) than those observed for R42 and site 49. Consistent with our experimental results showing a predictable change in glycation behavior at R54, we observed that there was a population shift towards shorter distances (≤ 9Å) for dialAGE^R54^(−), and a major population shift towards longer distances (> 9Å) for dialAGE^R54^(+) (Supporting Information, Figure S12, Figure S19). When we looked at the impact of the dialAGE^R54^ variants on the R42 site, we saw substantially more variability in the distances. However, the predicted trends did not match with our experimental results (Supporting Information, Figure S19). Together, these results suggest that dialAGE works better for sites at which mutations can be introduced within 5Å of the reactive Arg to avoid unexpected behaviors at other sites.

Next, to assess the impact of dialAGE on Ub function, we performed an *in vitro* auto-ubiquitination assay using recombinant E1 (Ube1), E2 (UbcH5a), and E3 (Mdm2) enzymes incubated with each dialAGE Ub variant. No apparent differences in the extent of polyubiquitination were observed, suggesting that the dialAGE mutations are non-perturbing to the ability of Ub to become conjugated to substrates. Next, we performed this same assay following a 24 h incubation with MGO, which was then quenched using Tris-HCl buffer (Supporting Information, Figure S20). Compared to their untreated counterparts, glycated dialAGE Ub exhibited comparable levels of polyubiquitination. We also found that this was the case when the concentration of MGO was doubled. These results suggest that Ub glycation does not interfere with its ability to become linked to substrate proteins *in vitro* (Supporting Information, Figure S21).

To date, there are no available methods that can control glycation of a specific protein in a cellular context, let alone at a specific site within that protein. Therefore, our next goal was to establish a proof-of-concept demonstration that dialAGE influences the site-specific glycation of Ub in living cells. To assess this, we expressed dialAGE^R42^ Ub variants fused to the N-terminus of an enhanced green fluorescent protein. Crucially, the fusion of GFP to the Ub C-terminus prevents the conjugation of dialAGE Ub-GFP to other substrate proteins, which would greatly diminish the signal of interest and complicate analysis. After confirming that all dialAGE^R42^ Ub-GFP variants were expressed comparably in HEK 293-T cells, glycation was induced by adding ± 5 mM MGO for 2 h prior to harvesting and lysis (Figure 4A,B). These samples were immunoprecipitated against GFP, and subsequently analyzed by mass spectrometry. Using intact protein LC-MS, we observed differences in AGE levels in the deconvoluted spectra (Figure 4C). This effect was most pronounced for the [M+54] adduct, which was the most clearly resolved AGE peak. Although the overall levels of glycation were lower than those observed *in vitro*, it was still possible to resolve differences in glycation at R42. Importantly, the AGE levels at R42 were modulated in the expected fashion ((+) > WT > (−)) (Figure 4E,G). The greatest extent of R42 glycation was observed for dialAGE^R42^(+) (8.5 ± 0.6%), and the least was observed for dialAGE^R42^(−) (4.7 ± 0.7%). However, we found that glycation at R54 (8.1 ± 1.7%, 9.1 ± 1.3%, and 7.1 ± 1.9%) and R72+R74 (12.7 ± 2.3%, 11.6 ± 2.8%, and 10.1 ± 2.3%) remained consistent across all dialAGE variants (Figure 4E-G). These conditions did not affect cell viability (Supporting Information, Figure S22), similar to a previous report.^[57]^ Due to the relatively low level of glycation, it was not possible to observe any glycation signal when using lower MGO concentrations. However, this effect was consistent when using higher (7 mM) MGO concentrations, resulting in increased glycation levels but reduced modulation between variants (Supporting Information, Figure S22). We also found that the use of dialAGE^R42^(−)^Q49D^ produced an even lower level of glycation at R42 than the original dialAGE^R42^(−), which improves the dynamic range of this approach (Figure S23). Taken together, these results demonstrate that dialAGE is promising new strategy for influencing site-specific glycation events in living cells.

**Figure 4.**
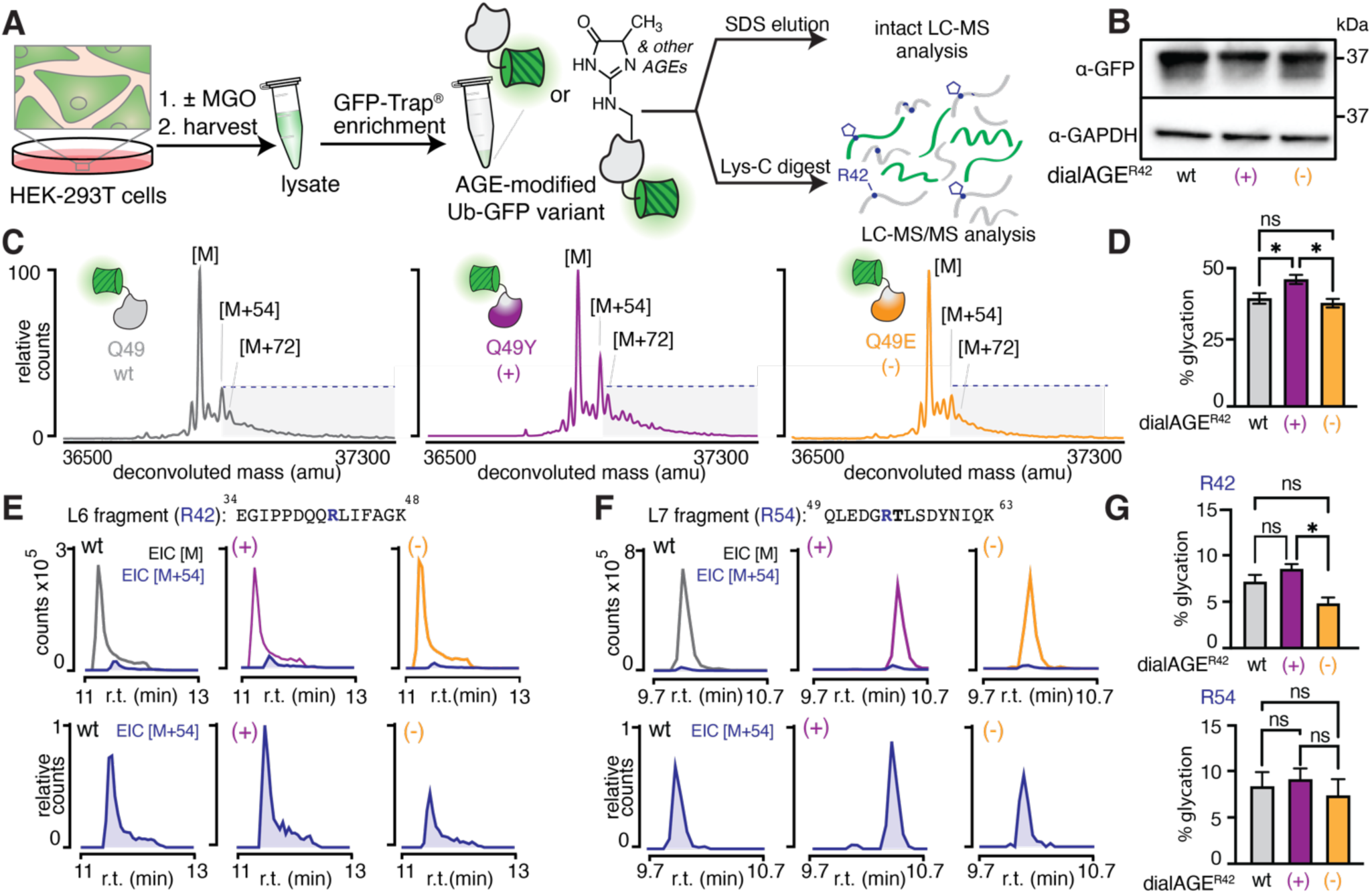
Influencing site-specific glycation of Ub in cells using dialAGE. A) HEK-293T cells expressing dialAGER42 versions of a Ub-GFP fusion protein were treated with or without 5 mM MGO for 2 h. Following enrichment against GFP, the extent of Ub-GFP glycation was determined using intact protein mass spectrometry or Lys-C digest followed by LC-MS/MS analysis. B) Western blot of HEK 293-T cell lysates probed for GFP and GAPDH show equivalent expression of all three dialAGE^R42^ variants. C) Intact mass spectra of dialAGE^R42^ Ub-GFP variants after treatment with MGO reveal observable differences in glycation. D) These differences were quantified by determining the intensity of [M+X] AGE peaks as a percent of total dialAGE^R42^ Ub-GFP. A nondirectional (two-tailed) one way ANOVA using Tukey’s multiple comparison test was used to determine statistically significant differences in glycation. P<.05 (*), p<.01 (**), p<.001 (***), p<.0001 (****). E) Extracted ion chromatograms (EIC) for the unmodified ([M]) or MGH-modified ([M+54]) R42 fragments show differences in glycation at R42. F) By contrast, EIC for the unmodified ([M]) or MGH-modified ([M+54]) R54 fragments show consistent levels of glycation at R42 across all three variants. Although the fusion of GFP introduced 7 additional Arg, we observed that glycation levels at those sites remained consistent for each dialAGE^R42^ variant. G) These findings were determined by quantifying the volume of AGE-modified fragments compared to the volume of all unmodified and modified fragments. Nondirectional (two-tailed) one-way ANOVA using Tukey’s multiple comparison test was used to determine statistically significant differences in glycation. P<.05 (*), p<.01 (**), p<.001 (***), p<.0001 (****).

## Conclusion

Here we have reported a new strategy, called dialAGE, that leverages our knowledge of the surrounding chemical microenvironment to tune glycation at a specific site.^[12,13]^ To our knowledge, this is the first method capable of modulating site-specific protein glycation events under identical concentrations of glycating agent. This technological advancement lays the groundwork for enabling tightly controlled biological experiments in which differences in glycation can be attained under uniform conditions, without changing the levels of glycating agent. Such an experimental set-up would enable direct comparisons that could unambiguously attribute a site-specific glycation event to a genuine biological function. For example, it has been reported that glycation impairs proteasomal degradation^[58,59]^ and/or is cleared by autophagy, but there are no studies that have evaluated how site-specific Ub glycation influences the function of Ub or the UPS.^[43–47,60]^ For this reason, we expect that this tool will be useful not only for elucidating the consequences of Ub glycation in the context of proteasomal degradation, but also for investigating the impacts of glycation on the numerous other disease-relevant proteins that have had reported alterations in their *in vitro* activities following glycation.^[6,43–45,61–64]^ Our future work will pursue both of these exciting new avenues for further study.

In this study, we relied on chemical insights generated from our prior work, which revealed key sequence features like Tyr or Glu as major drivers of glycation susceptibility.^[12,13]^ Our initial development of dialAGE focused solely on these most apparently influential sequence elements, which we found to broadly enhance or suppress overall glycation levels, respectively. Our results imply that the strength of dialAGE-dependent glycation modulation is sensitive to distance, which is consistent with our prior findings in the context of short, unstructured peptides. On proteins, the introduced mutations resulted in the expected modulation of overall Ub glycation levels for both dialAGE^R42^ and dialAGE^R54^, but we also found that only dialAGE^R42^ produced a consistently site-specific effect. Because we ruled out the possibilities that this could be attributed to a major disruption of the Ub fold or from differences in concentration, we therefore conclude that any dialAGE variant must be carefully vetted prior to use. Additionally, molecular dynamics simulations proved highly useful in understanding how dialAGE works. In future studies, we plan to introduce MGO molecules in the simulations of Ub variants to understand how various mutations affect MGO–Ub interactions, which will provide further insights into dialAGE performance. Taken together, these results also suggest that it would be possible to use dialAGE to broadly modulate glycation at all sites within a protein, not just one, which may be beneficial—and perhaps even preferred—for certain applications. For these reasons, our future work will focus on expanding the repertoire of side chain functionalities that produce a predictable dialAGE effect. In particular, a deeper understanding of residues that influence AGE distributions—not only total AGE levels—will be of great benefit for determining the biological functions of specific AGEs.

## Supporting Information

The authors have cited additional references within the Supporting Information.^[65–70]^

## Acknowledgements

This work was supported by National Institutes of Health grant R01GM132422 to R.A.S, as well as by a gift to the Scheck Lab from J. Kanagy, and a gift to the Scheck Lab from J. Fickel. The authors thank J. McEwen for helpful comments on the manuscript.

## Notes

### Competing Interest Statement

The authors have declared no competing interest.

