## Supporting Information for "Predictable modulation of a spontaneous post-translational modification in living cells"

Meghan S. Martin, Nomindari Bayaraa, Brittany T. Fox, Yu-Shan Lin, and Rebecca A. Scheck*
Department of Chemistry, Tufts University, 62 Talbot Avenue, Medford, Massachusetts 02155

### Table of Contents

#### Materials & Methods


| | | |
|---|---|---|
| Experimental Procedures | | 1 |


#### Supporting Figures and Tables


| | | |
|---|---|---|
| **Figure S1** | Purification and Characterization of dialAGE$^{R42}$ Ub variants | 8 |
| **Figure S2** | Comparison of the reactivity of glucose and MGO | 9 |
| **Figure S3** | Optimization of MGO treatment conditions for HA-Ub | 10 |
| **Figure S4** | Total glycation levels are modulated with dialAGE | 11 |
| **Figure S5** | DialAGE$^{R42}$ leads to site-specific differences in R42 glycation, as revealed by Lys-C digest | 12 |
| **Figure S6** | DialAGE$^{R42}$ R72+74 display minor differences in glycation | 13 |
| **Figure S7** | DialAGE$^{R42}$ leads to site-specific differences in R42 glycation, as revealed with trypsin digest | 14 |
| **Figure S8** | DialAGE$^{WT}$ and commercially HA-Ub are similarly glycated | 15 |
| **Figure S9** | DialAGE influences glycation rates | 16 |
| **Figure S10** | DialAGE does not significantly modulate AGE distributions | 17 |
| **Figure S11** | Gln mutation to Asp leads to amplified decrease in R42 glycation | 18 |
| **Figure S12** | Trends in root-mean-square deviation (RMSD), root-mean-square fluctuation (RMSF), radius of gyration, and solvent accessible surface area (SASA) cannot explain experimentally observed dialAGE behavior | 19 |
| **Figure S13** | DialAGE$^{R42}$ modulates distances surrounding R42 but not R54 | 20 |
| **Figure S14** | Purification and characterization of dialAGE$^{R54}$ Ub variants | 21 |
| **Figure S15** | Glycation modulation at R54 using dialAGE$^{R54}$ | 22 |
| **Figure S16** | DialAGE$^{R54}$ modulates glycation levels but not AGE distributions | 23 |
| **Figure S17** | Circular dichroism at elevated temperature reveals no structural differences in dialAGE$^{R54}$ variants | 24 |
| **Figure S18** | Phenylalanine mutation at T55 does not impact glycation | 25 |
| **Figure S19** | Molecular dynamics simulations for dialAGE$^{R54}$ | 26 |
| **Figure S20** | MGO treatment does not change protein concentrations | 27 |
| **Figure S21** | *In vitro* ubiquitination using dialAGE Ub variants | 28 |
| **Figure S22** | Ub-GFP dialAGE$^{R42}$ variants have scaled levels of glycation with increasing MGO | 29 |
| **Figure S23** | Ub-GFP dialAGE$^{R42}$(-)$^{Q49D}$ leads to greater dynamic range in glycation signal | 30 |
| **Table S1** | Ub DialAGE Lys-C Fragments | 31 |
| **Table S2** | Ub DialAGE Tryptic Fragments | 32 |
| **Table S3** | Expected and observed fragments for R42 | 33 |
| **Table S4** | Expected and observed fragments for R54 | 34 |
| **Table S5** | Expected and observed fragments for R72+74 | 35 |

| | | |
|---|---|---|
| Supporting References | | 36 |

### Experimental Procedures

#### General Materials

All chemical reagents and solvents were of analytical grade, obtained from commercial suppliers and used without further purification unless otherwise noted. Methylglyoxal (40% w/v in water) was purchased from MilliporeSigma (M0252). Commercial HA-Ub was purchased from R&D systems (U-110).

#### Cloning

#### dialAGE$^{WT}$ Ub sequence

Orange bases denote the N-terminal 6x-His and HA epitopes, while black bases are ubiquitin derived. Red bases represent the codon for Q49 and T55.

```
ATGTCGTACTACCATCACCATCACCATCACCTCGAATCAACAAGTTTGTACAAAAAAGCAGGCT
TCACCATGGGCAGCATGTACCCCTACGACGTGCCAGATTACGCCAGCGGCTCCATGCAGATCTT
CGTGAAAACCCTGACCGGCAAGACCATCACCCTGGAAGTGGAACCCAGCGACACCATCGAGAAT
GTGAAAGCCAAGATCCAGGACAAAGAGGGCATCCCCCCCGACCAGCAGAGACTGATCTTCGCCG
GAAAGCAGCTGGAAGATGGCCGGACCCTGAGCGACTACAACATCCAGAAAGAGTCCACCCTGCA
CCTGGTGCTGAGGCTGAGAGGCGGATGA
```

#### Ub-GFP sequence

Black bases are ubiquitin derived, while green bases correspond to GFP and linker bases. Red bases represent the codon for Q49.

```
ATGCAGATCTTCGTGAAGACTCTGACTGGTAAGACCATCACCCTCGAGGTTGAGCCCAGTGACA
CCATTGAGAATGTCAAGGCAAAGATCCAAGATAAGGAAGGCATCCCTCCTGACCAGCAGAGGCT
GATCTTTGCTGGAAAACAGCTGGAAGATGGGCGCACCCTGTCTGACTACAACATCCAGAAAGAG
TCCACCCTGCACCTGGTACTCCGTCTCAGAGGTGTGGTGGGGAAGCTTGGTCGACAGGATCCAC
CGGTCGCCACCATGGTGAGCAAGGGCGAGGAGCTGTTCACCGGGGTGGTGCCCATCCTGGTCGA
GCTGGACGGCGACGTAAACGGCCACAAGTTCAGCGTGTCCGGCGAGGGCGAGGGCGATGCCACC
TACGGCAAGCTGACCCTGAAGTTCATCTGCACCACCGGCAAGCTGCCCGTGCCCTGGCCCACCC
TCGTGACCACCCTGACCTACGGCGTGCAGTGCTTCAGCCGCTACCCCGACCACATGAAGCAGCA
CGACTTCTTCAAGTCCGCCATGCCCGAAGGCTACGTCCAGGAGCGCACCATCTTCTTCAAGGAC
GACGGCAACTACAAGACCCGCGCCGAGGTGAAGTTCGAGGGCGACACCCTGGTGAACCGCATCG
AGCTGAAGGGCATCGACTTCAAGGAGGACGGCAACATCCTGGGGCACAAGCTGGAGTACAACTA
CAACAGCCACAACGTCTATATCATGGCCGACAAGCAGAAGAACGGCATCAAGGTGAACTTCAAG
ATCCGCCACAACATCGAGGACGGCAGCGTGCAGCTCGCCGACCACTACCAGCAGAACACCCCCA
TCGGCGACGGCCCCGTGCTGCTGCCCGACAACCACTACCTGAGCACCCAGTCCGCCCTGAGCAA
AGACCCCAACGAGAAGCGCGATCACATGGTCCTGCTGGAGTTCGTGACCGCCGCCGGGATCACT
CTCGGCATGGACGAGCTGTACAAGTAA
```

**Primers used**
A pDEST17 (Invitrogen) plasmid containing the dialAGE$^{WT}$ Ub sequence was used as a template for site-directed mutagenesis. All dialAGE variants were prepared using a QuikChange XL II kit (Agilent) with the following 5'-3' primers:

| | | |
|---|---|---|
| dialAGE$^{R42}$(+) | Q49Y forward | TCCGGCCATCTTCCAGATACTTTCCGGCGAAGATC |
| dialAGE$^{R42}$(+) | Q49Y reverse | GATCTTCGCCGGAAAGTATCTGGAAGATGGCCGGA |
| dialAGE$^{R42}$(-) | Q49E forward | GCCATCTTCCAGCTCCTTTCCGGCGAAGA |
| dialAGE$^{R42}$(-) | Q49E reverse | TCTTCGCCGGAAAGGAGCTGGAAGATGGC |
| dialAGE$^{R54}$(-) | T55E forward | GATGTTGTAGTCGCTCAGTTCCCGGCCATCTTCCAGCTG |
| dialAGE$^{R54}$(-) | T55E reverse | CAGCTGGAAGATGGCCGGGAACTGAGCGACTACAACATC |
| dialAGE$^{R54}$(+) | T55Y forward | GTTGTAGTCGCTCAGGTACCGGCCATCTTCCAGC |
| dialAGE$^{R54}$(+) | T55Y reverse | GCTGGAAGATGGCCGGTACCTGAGCGACTACAAC |
| dialAGE$^{Q49D}$ | Q49D forward | CATCTTCCAGGTCCTTTCCGGCGAAGATCAG |
| dialAGE$^{Q49D}$ | Q49D reverse | CTGATCTTCGCCGGAAAGGACCTGGAAGATG |
| dialAGE$^{Q49S}$ | Q49S forward | CATCTTCCAGACTCTTTCCGGCGAAGATCAG |
| dialAGE$^{Q49S}$ | Q49S reverse | CTGATCTTCGCCGGAAAGAGTCTGGAAGATG |
| dialAGE$^{T55F}$ | T55F forward | GTTGTAGTCGCTCAGGAACCGGCCATCTTCCAGC |
| dialAGE$^{T55F}$ | T55F reverse | GCTGGAAGATGGCCGGTTCCTGAGCGACTACAAC |

Ub-GFP G76V plasmid was obtained from Nico Dantuma (Addgene #11941). Q49E and Q49Y variants were cloned via QuikChange XL II kit with the following 5'-3' primers:

| | | |
|---|---|---|
| dialAGE$^{R42}$(+) | Q49Y forward | CATCTTCCAGATATTTTCCAGCAAAGATCAG |
| dialAGE$^{R42}$(+) | Q49Y reverse | CTGATCTTTGCTGGAAAATATCTGGAAGATG |
| dialAGE$^{R42}$(-) | Q49E forward | CATCTTCCAGCTCTTTTCCAGCAAAGATCAG |
| dialAGE$^{R42}$(-) | Q49E reverse | CTGATCTTTGCTGGAAAAGAGCTGGAAGATG |

| | | |
|---|---|---|
| dialAGE$^{R42}$(-)$^{Q49D}$ | Q49D forward | CATCTTCCAGGTCTTTTCCAGCAAAGATCAG |
| dialAGE$^{R42}$(-)$^{Q49D}$ | Q49D reverse | CTGATCTTTGCTGGAAAAGACCTGGAAGATG |

**Protein Expression and Purification**

To express dialAGE Ub variants for use *in vitro*, BL21 (DE3) (Agilent) *E. coli* were transformed with the indicated plasmids. After starter cultures (5 mL) were incubated at 37 ºC overnight in LB with continuous shaking, they were expanded into 100 mL cultures and incubated overnight, then diluted into 1 L cultures with continuous shaking until reaching $OD_{600}$ = 0.45–0.6. At this time, 0.4 mM isopropyl β-D-1-Thiogalactopyranoside (IPTG) was added to induce expression. After 5 h incubation at 18 ºC, the resulting cells were pelleted and frozen. Lysis was accomplished using sonication on ice for 3.5 min in 50 mM Tris-HCl pH 7.0, 500 mM NaCl, 1 mM MgCl, with 40 U Benzonase (MilliporeSigma), and 0.4 mM phenylmethylsulfonyl fluoride. The resulting lysates were purified on a Ni-NTA column (Cytiva His GraviTrap™), washing with 20 mM sodium phosphate pH 7.4, 20 mM imidazole, 500 mM NaCl. After elution with a buffer containing 20 mM sodium phosphate pH 7.4, 500 mM NaCl with 500 mM imidazole, the resulting purified dialAGE Ub variants were buffer exchanged into 20 mM PBS at pH 7.4 (Cytiva Nap™-25 column). Protein concentration was measured via Bradford Assay (Pierce) and pure proteins were aliquoted and stored at -20 ºC.

**Circular Dichroism**

Samples were prepared at 15 µM concentration in 20 mM phosphate buffer at pH 7.4. CD spectra were acquired on a Jasco J-815 CD Spectrometer in quartz cells of 0.1 cm path length using the following parameters: wavelength, 190-260 nm; step resolution, 0.5 nm; speed, 20 nm/min; accumulations, 3; response, 1 sec; bandwidth, 0.5 nm. Using the spectrometer software, background noise was removed using the appropriate blank control and spectra were smoothed.

**General Protocol for in vitro Protein Glycation.**

To perform *in vitro* glycation, dialAGE variants (final concentration 50 µM) were incubated with the desired concentration of methylglyoxal (MGO, 100 or 200 µM, 2 or 4 equivalents) in 20 mM PBS, pH 7.4 for 24 h (and/or the indicated time, up to 48 h) at 37 °C without agitation. Following reaction, samples were quenched with a final concentration of 5 mM Tris-HCl pH 7.3, diluted with ultrapure water and subsequently subjected to LC-MS analysis.

**Proteolytic Digestion**

For digestion of proteins after MGO treatment, the resulting protein solutions were diluted with additional 1:1 t-butanol:water and DTT was added to a final concentration of 3 mM, with 0.5 µL of 1% Trypsin Enhancer (Promega). Reactions were heated to 65°C for 35 minutes and subsequently cooled to room temperature. Sequencing grade modified trypsin (Promega) or Lys-C (Mass Spec grade, Promega) was added for a final 1:25 protease:protein ratio in 40 µL total volume, along with an additional 0.5 µL of Trypsin

Enhancer. The digest solution was incubated at 37 °C for 5 h. The resulting solution of peptides was subjected to LC-MS/MS analysis without further purification.

**Liquid Chromatography-Mass Spectrometry Acquisition and Analysis.**
Reversed-phase chromatography and mass spectrometry were performed on an Agilent 1260 Infinity LC system in line with an Agilent 6530 Q-TOF. Undigested protein samples were diluted in water and injected onto a Zorbax 300 SB-C8 Rapid Resolution HD 1.8 µm column (2.1 x 100 mm, Agilent), eluting with a water:acetonitrile gradient mobile phase with 0.1% formic acid (0.400 mL/min; 95% - 20% water over 26 min). The mass spectrometer was utilized in positive mode with a dual electrospray ionization (ESI) source. MS spectra were acquired using the following settings: ESI capillary voltage, 4500 V; fragmentor, 250 V; gas temperature, 325°C; gas rate, 12.5 L/min; nebulizer, 50 psig. Data was acquired at rate of 5 spectra per second and scan range of 100–3000 m/z. Analysis was accomplished with Agilent MassHunter BioConfirm (v10.0) using the Intact Protein workflow.

**Equation S1**

$$\%\,unmodified\ protein = \left(\frac{volume\ unmodified\ protein\ [M]}{total\ volume\ modified\ and\ unmodified\ protein}\right) * 100$$

Following tryptic digestion, peptide fragments were injected onto an AdvanceBio Peptide 2.7 µm column (2.1 x 150 mm, Agilent) and were eluted with a water:acetonitrile gradient mobile phase with 0.1% formic acid (0.400 mL/min; 95% - 5% water over 19 min). MS spectra were acquired using the following settings: ESI capillary voltage, 4000 V; fragmenter, 150 V; gas temperature, 325°C; gas rate, 12 L/min; nebulizer, 40 psig. Data was acquired at a rate = 5 spectra per second and scan range of 300–3000 m/z. MS/MS spectra were acquired using the following settings: ESI capillary voltage, 4000 V; fragmentor, 150 V; gas temperature, 325°C; gas rate 12 L/min; nebulizer, 40 psig. MS/MS was acquired at 2 spectra per second with a mass range of 100–3000 m/z, with stringency set to a medium isolation width. After identification, precursor ions were subjected to iterative rounds of collision induced dissociation in the collision chamber and subsequent mass identification. A ramped collision energy was used with a slope of 3.6 and offset of -4.8 as well as a slope of 3 and offset of 2. MS data was quantified using the MassHunter Molecular Feature Extractor in MassHunter Qualitative Analysis (v. 10.00), which reports cumulative MS ion counts as ‘volumes’ observed for any and all charge states associated with a particular compound. Quantification (% modification) was carried out by dividing the specified mass adduct(s) volume by the total volume of modified and unmodified peptide.

**Equation S2**

$$\%\,modification = \left(\frac{volume\ modified\ peptide}{volume\ modified + unmodified\ peptide}\right) * 100$$

**Molecular dynamics (MD) simulations**
The initial conformation of ubiquitin (Ub) was extracted from the PDB entry 1UBQ.[1] We employed the Chimera[2] software package to generate conformations of dialAGE$^{WT}$, dialAGE$^{R42}$(+) (Q49Y), dialAGE$^{R42}$(-) (Q49E), dialAGE$^{R54}$(+) (T55Y), dialAGE$^{R54}$(-) (T55E)

Ub variants. All proteins were prepared with standard protonation states corresponding to the physiological pH of 7. Because the experiments employed Ub variants with N-terminal 6×-His and HA epitope tags, the N-terminus of each Ub variant was capped by an acetyl group, and the C-terminus was left uncapped as a free carboxyl group in the simulations.

All MD simulations were performed with GROMACS 2023[3] using the Amber ff19SB[4] force field with the OPC water model.[5] We chose the ff19SB force field because it was the most up-to-date Amber protein force field and was benchmarked on 1UBQ.[4] For each system, the initial structure was solvated in a box of pre-equilibrated water molecules, with the minimum distance between the protein and the walls of the box of at least 1.0 nm. When the total charge of the protein was nonzero, minimal simple counterions ($Na^+$) were added to neutralize the whole system. The solvated system was energy-minimized with the steepest descent algorithm. During the energy minimization, all protein heavy atoms were positionally restrained by a harmonic potential with a force constant of 1,000 kJ $mol^{-1}$ $nm^{-2}$. The same positional restraints were maintained as a 10-ns NVT simulation at 300 K was conducted. During this NVT step, 5 parallel runs with different initial velocities were generated for each system. After the NVT simulation, the same positional restraints were maintained for a subsequent 10-ns NPT simulation at 300 K and 1 bar. To allow the sidechains to relax, positional restraint with a force constant of 1,000 kJ $mol^{-1}$ $nm^{-2}$ was applied only to the backbone heavy atoms for a subsequent 10-ns NVT simulation at 300 K followed by a 10-ns NPT simulation at 300 K and 1 bar. Then all positional restraints were removed to allow the whole system to equilibrate. The non-constrained system was equilibrated again for a 10-ns NVT simulation at 300 K and a 10-ns NPT simulation at 300 K and 1 bar. Finally, the production NPT simulation was carried out for 200 ns at 300 K and 1 bar. Dynamics evolution utilized the leapfrog algorithm with a time step of 2 fs. Temperature control was achieved with v-rescale thermostat separately regulating the protein and solvent with coupling time constants of 0.1 ps. Pressure control was achieved with Parrinello-Rahman barostat with a coupling time constant of 2 ps and an isothermal compressibility of $4.5 \times 10^{-5}$ $bar^{-1}$. Bonds involving hydrogen were constrained with LINCS algorithm. Electrostatic and van der Waals interactions were truncated at 1.0 nm. Long-range electrostatics were treated using the particle mesh Ewald summation with a Fourier grid spacing of 0.12 nm and an order of 4. A long-range dispersion correction was applied to correct energy and pressure. The last 100 ns of the production run was used for subsequent analysis.

***In vitro* auto-ubiquitination assay**
To assess the ability of dialAGE Ub variants to be incorporated into polyubiquitinated chains, 5 µM of each dialAGE variant (treated with or without MGO, as described above) was incubated for 1 h at 37° C with 500 nM E1 enzyme (His6-Ube1, R&D systems), 1 µM E2 enzyme (UbE2D1, South Bay Bio), and 1 µM E3 enzyme (GST-Hdm2 RING domain, Enzo Life Sciences), along with 2 mM Mg-ATP, and 1 mM DTT in 40 mM Tris, pH 7.6. After 1 h, autoubiquitination reactions were quenched by addition of 10X SDS to a final concentration of 2X SDS. 2 µL of loading dye+DTT was added to 10 µl of the reaction dilution, boiled at 95° C, spun in a mini centrifuge, and the resulting supernatant was loaded and run on a 4–15% gradient precast SDS-PAGE gel (mini-PROTEAN

TGX™) in standard Tris/glycine/SDS running buffer at 226 mV for 26 min to resolve protein bands. Following separation by SDS-PAGE, proteins were transferred to PVDF membrane using the iBlot 2 (Invitrogen) dry blotting system. Membranes were subsequently blocked in a buffer containing 20 mM Tris, 150 mM NaCl and 0.1% tween (1X TBST) supplemented with 5% (w/v) Bovine serum albumin (BSA). After blocking, primary antibody incubation was achieved overnight at 4 °C. Membranes were washed 3X for 5 min each with 1X TBST. Detection was performed by incubating membranes with HRP-linked secondary antibodies (Cell Signaling Technology) in 5% BSA-TBST for one hour at room temperature. After secondary antibody incubation, membranes were washed again 3X for 5 min with 1X TBST. Chemiluminescent signal was developed with Clarity Western ECL Substrate (Bio-Rad) and imaged on a Bio-Rad ChemiDoc XRS+. The following antibodies were used:

| Antibody | Vendor (product number) | Host species | dilution |
|---|---|---|---|
| α-HA | Cell Signaling Technologies (2367) | mouse | 1:1000 |
| α-mouse-HRP | Cell Signaling Technologies (7076) | | 1:2000 |

**Mammalian Cell Culture**

HEK-293T cells were cultured under standard conditions in Dulbecco's Modified Eagle Medium (DMEM) supplemented with 10% Fetal Bovine Serum (FBS) and 1% Penicillin-Streptomycin (pen/strep) at 37 °C at 5% $CO_2$. Cells were passaged every 2 to 3 days (80–90% confluency) for 25-30 passages.

**General Protocol for Cellular Glycation.**

Roughly 1,000,000 HEK 293-T cells were seeded in a 60 mm dish and grown to ~60% confluency over ~18 h. DialAGE Ub-GFP variants were overexpressed by transfecting cells with 3 µg of the desired Ub-GFP dialAGE plasmid using TransIT-LT1 Transfection Reagent (Mirus Bio, 6 µL/µg plasmid). After transfection, cells were grown for another 24 h to about 90% confluency. Media was removed and replaced with new media containing either 0 or 5 mM MGO for 2 hr. After treatment, cells were harvested with TrypLE Express (Gibco) dissociation reagent. Cells were lysed on ice with 500 µL Tris-Cl buffer (50 mM Tris, 150 mM NaCl, 1 mM EDTA, 1 mM NaF, 1% Triton X-100) at pH 7.5 with a Pierce Protease and Phosphatase Inhibitor tablet (1 tablet/10 mL buffer). Lysates were clarified by centrifugation at 4255 x g for 30 min, and total protein quantified by BCA Protein Assay (Pierce) on a Tecan Spark 10M plate reader.

**Immunoprecipitation Protocol**

After clarification (described above), 500 µg protein of resulting cellular lysates were diluted to a total volume of 500 µL with 10 mM Tris, 150 mM NaCl, 0.5 mM EDTA, pH 7.5. To this solution, 30 µL of GFP-Trap® magnetic agarose slurry was added and allowed to incubate for overnight at 4° C with end-over-end rotation. Following the incubation period, the resin was pelleted and washed 3X with the same buffer. dialAGE Ub-GFP fusion proteins were either digested with Lys-C on bead, or full-length proteins were eluted by resuspending the protein-bound GFP-Trap® agarose in 50 µL of 2x SDS elution buffer and heating to 95° C for 5 min. Resulting eluates were analyzed by LC-MS/MS.

**Trypan Blue Analysis**

To measure cell viability of cells treated with either 0 or 5 mM glucose, following harvesting, cells were mixed 1:1 with 0.4% trypan blue solution (Gibco #15250061) and viability was measured on a Tecan Spark 10M plate reader.

### Supporting Figures and Tables

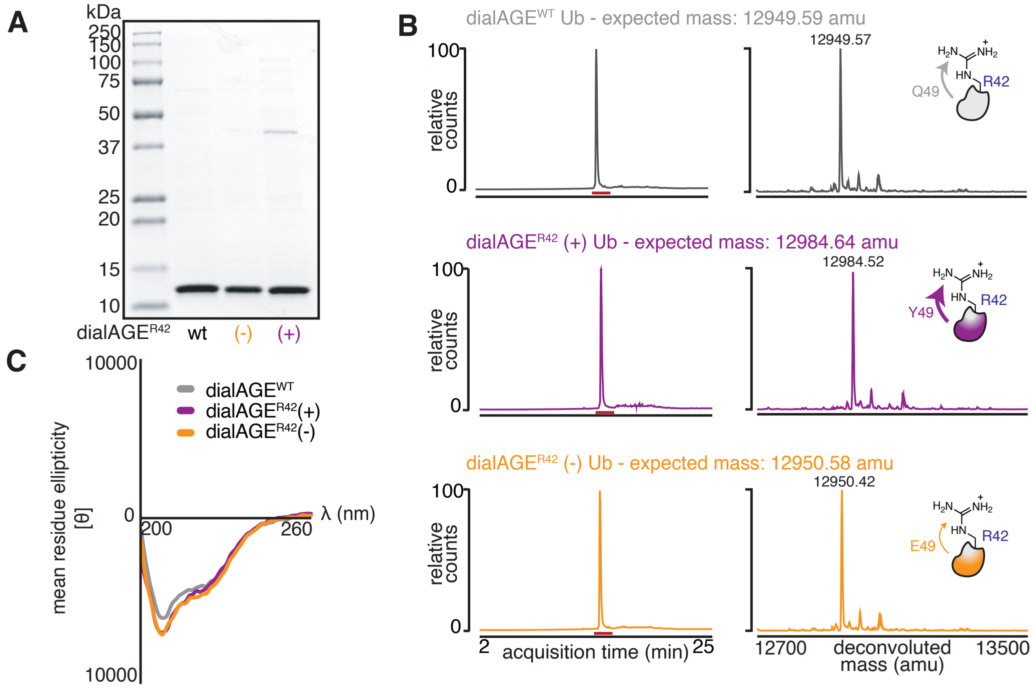


**Figure S1. Purification and Characterization of dialAGE$^{R42}$ Ub variants.** Following expression and purification, dialAGE$^{R42}$ Ub variants were assessed by **A)** Coomassie and **B**) LC-MS, revealing >95% purity for all variants. **C**) To evaluate dialAGE Ub structure, circular dichroism spectra were obtained for all variants, revealing nearly identical folds. The resulting CD spectra are very similar to published CD spectra of Ub, though small differences [6–8] are attributed to the addition of both an HA epitope and a 6x-His tag in dialAGE variants. Published Ub alanine scan spectra[7] show that mutation at even a single site can produce CD spectra with large differences; a lack of such difference in these spectra further supports that dialAGE Ub mutations do not grossly perturb Ub structure.

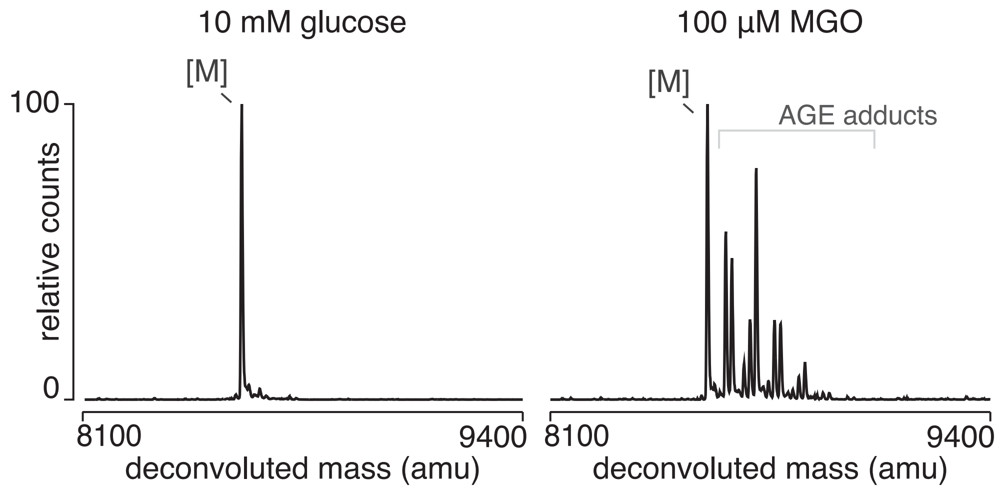


**Figure S2. Comparison of the reactivity of glucose and MGO**. Following incubation of ubiquitin with either 10 mM glucose (left) or 100 µM MGO (right) for 24 h, the extent of modification was observed. No apparent glycation was observed in glucose treated ubiquitin, while MGO treatment led to a robust amount of AGE adducts formed, confirming MGO to be a more reactive, and relevant, glycating agent than glucose.

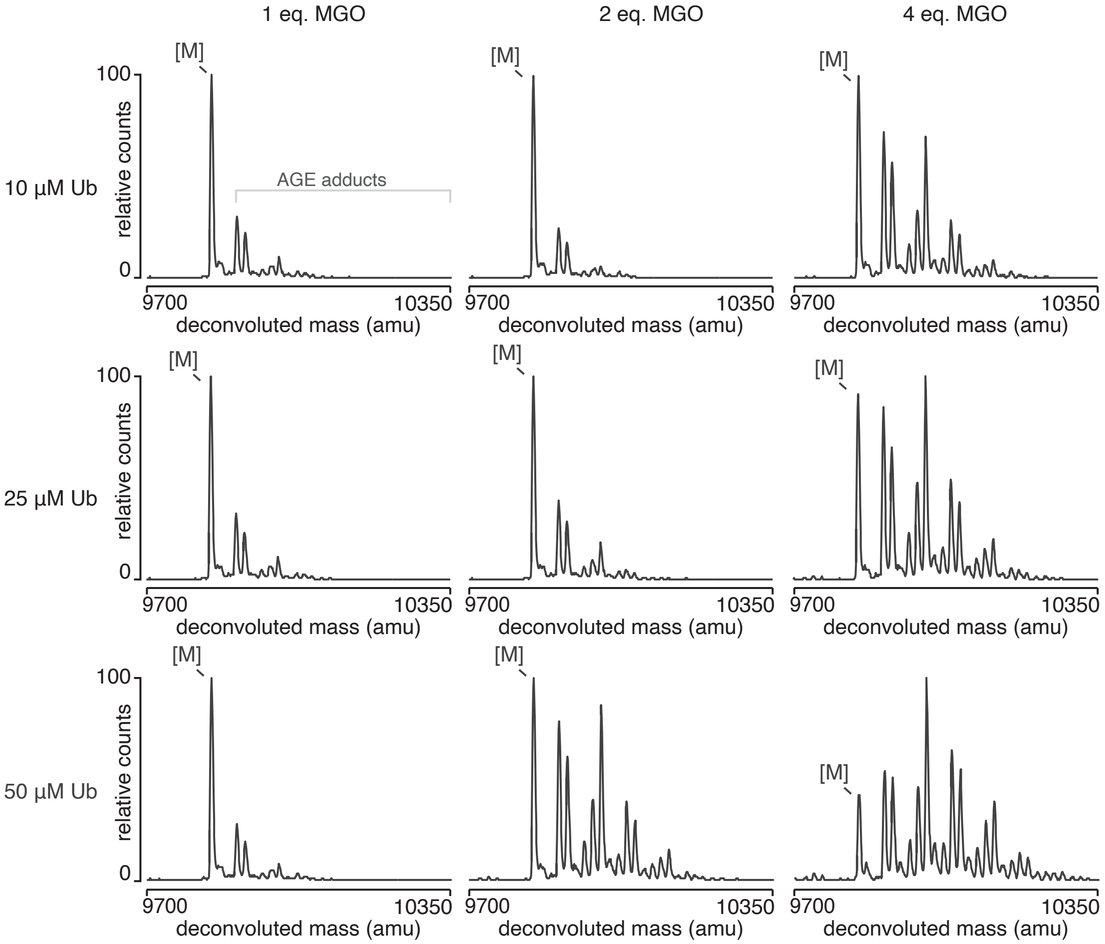


**Figure S3. Optimization of MGO treatment conditions for HA-Ub.** Commercially available HA-Ub was incubated at several different concentrations (10–50 µM) using different ratios of MGO (10–200 µM, 1–4 equiv.). These results revealed 50 µM HA-Ub treated with 2 equivalents MGO for 24 h in 20 mM PBS pH 7.4 was an optimal concentration, enabling clear observation of both increases and decreases in glycation.

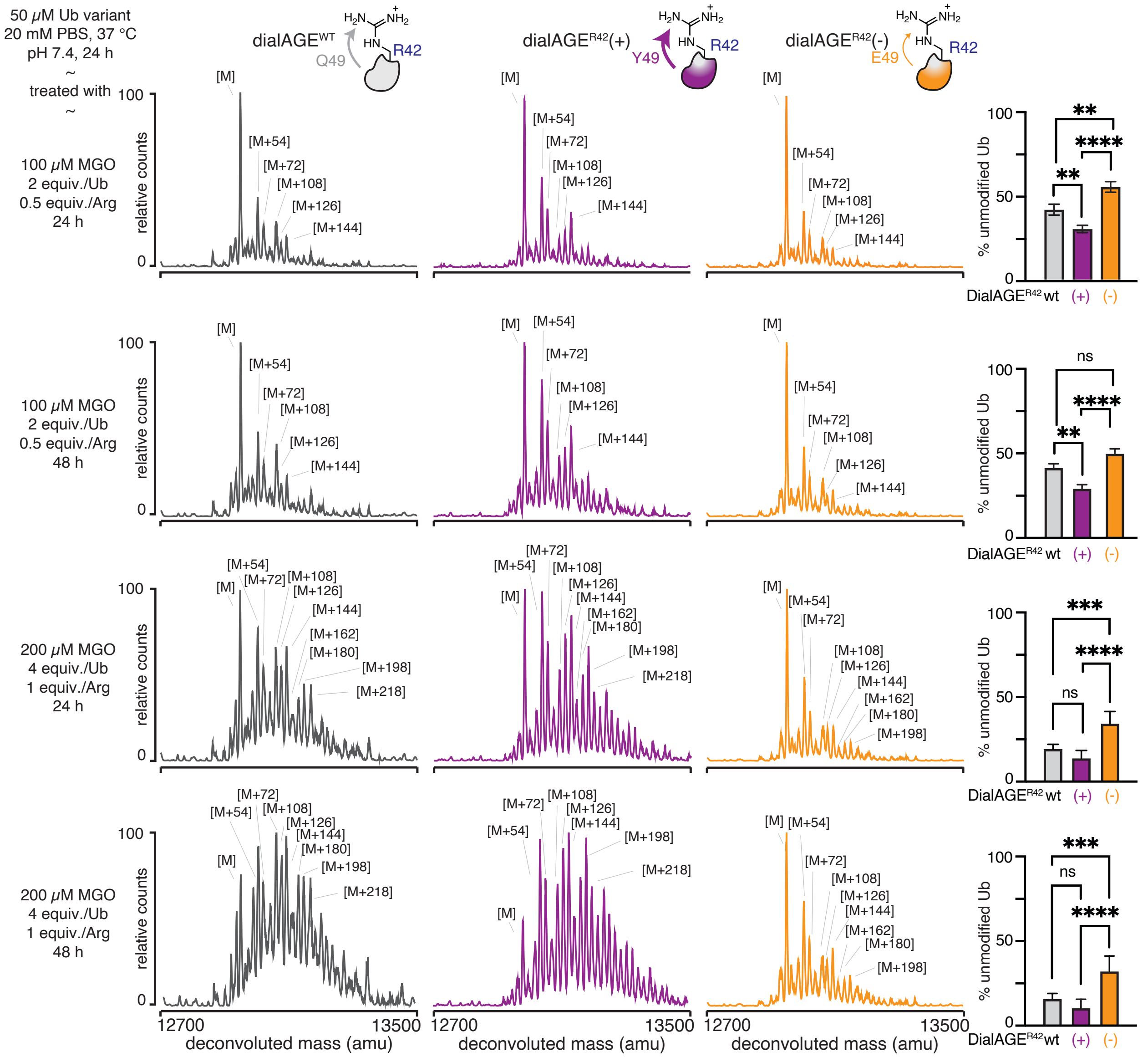


**Figure S4. Total glycation levels are modulated with dialAGE.** Recombinant dialAGE$^{WT}$ and dialAGE$^{R42}$ variants (50 μM) were treated with either 2 or 4 equivalents of MGO (100 or 200 μM) for 24 or 48 h and analyzed by intact protein mass spectrometry. Total levels of glycation were determined by quantifying the intensity of the remaining unmodified peak ([M]) after MGO treatment. In all cases, clear dialAGE-dependent differences in the total level of glycation were observed, with dialAGE$^{R42}$(+) leading to the most glycation, followed by dialAGE$^{WT}$, which was followed by dialAGE$^{R42}$(-). Bar graphs represent mean ± SEM. Nondirectional (two-tailed) two-way ANOVA using Tukey's multiple comparison tests were used to determine if each variant yielded statistically significant differences in glycation. P<.05 (*), p<.01 (**), p<.001 (***), p<.0001 (****).

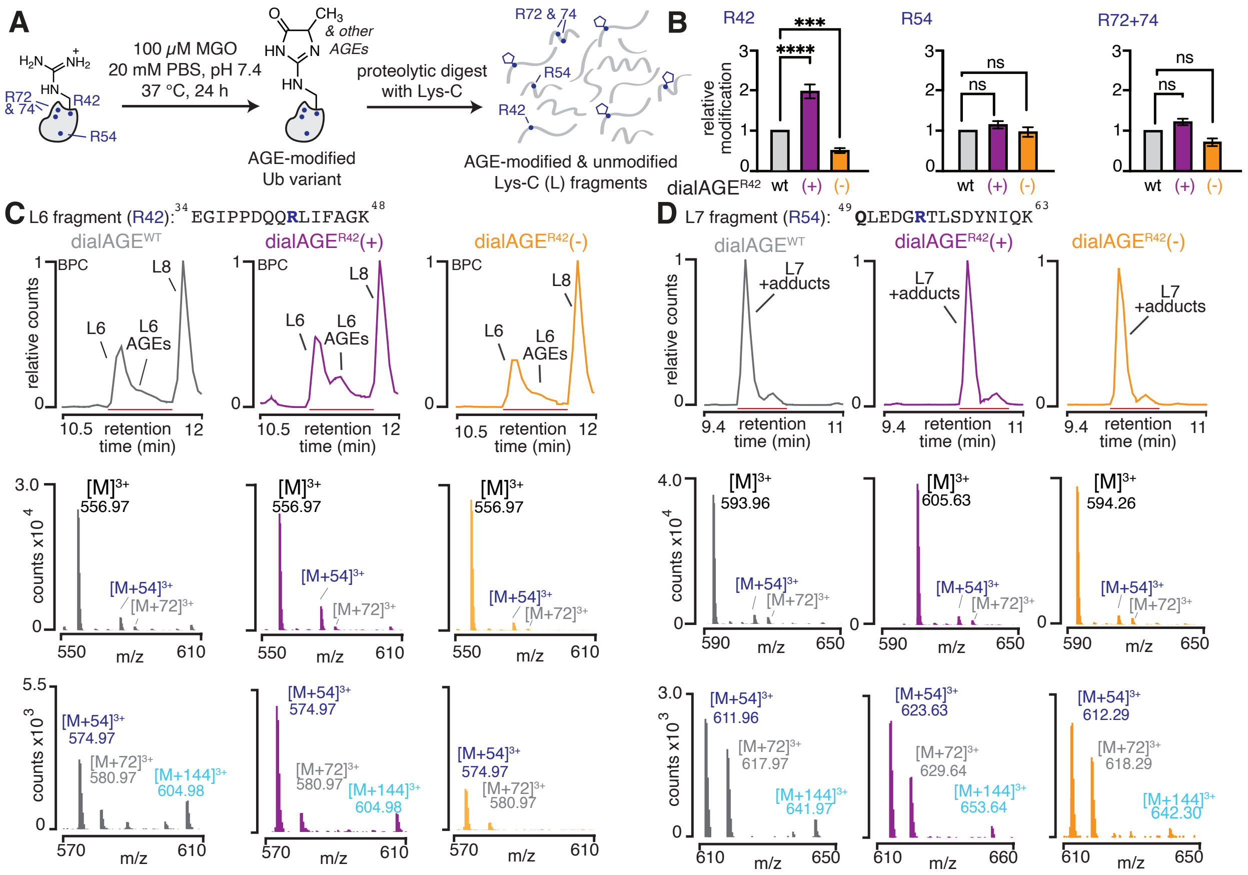


**Figure S5. DialAGE$^{R42}$ leads to site-specific differences in R42 glycation, as revealed by Lys-C digest. A)** dialAGE$^{WT}$ and dialAGE$^{R42}$ variants (50 µM) were treated with 2 equiv. MGO (100 µM) and subsequently digested by Lys-C. **B)** Quantification of relative glycation levels for each Arg reveal statistically significant differences solely at R42 for dialAGE$^{R42}$ Ub variants. **C)** Base peak chromatograms (BPC) (*top*) and combined MS spectra at the highlighted retention time showing *m/z* for unmodified and glycated R42 (L6 peptides) (*middle*), including zoomed in spectra that more clearly show *m/z* for modified R42 fragments (*bottom*). **D)** BPC (*top*) and combined MS spectra at the highlighted retention time showing *m/z* for unmodified and glycated R54 (L7 peptides) (*middle*), including zoomed in spectra that more clearly show *m/z* for modified R54 fragments (*bottom*). Together, these results suggest that dialAGE influences overall glycation levels by modulating AGE levels exclusively at a specific site. Bar graphs represent mean ± SEM. Nondirectional (two-tailed) two-way ANOVA using Tukey's multiple comparison tests were used to determine if each variant yielded statistically significant differences in glycation. P<0.05 (*), p<0.01 (**), p<0.001 (***), p<0.0001 (****).

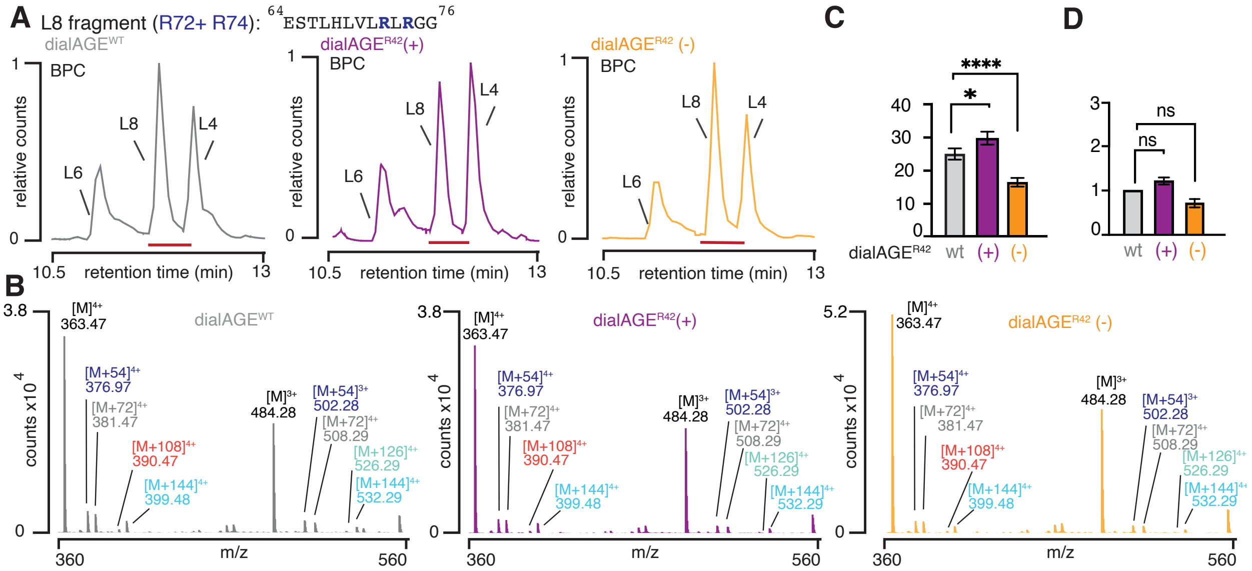


**Figure S6. DialAGE$^{R42}$ R72+74 display minor differences in glycation A)** Base peak chromatograms (BPC) for the dialAGE$^{R42}$ Lys-C fragment containing R72+R74 Lys-C fragments retention time. **B)** Combined spectra at the highlighted retention time show similar modification of R72+74 fragments between dialAGE$^{R42}$(+), dialAGE$^{WT}$ and dialAGE$^{R42}$(-). **C)** Absolute % glycation and **D)** relative glycation following 24 h treatment with 2 eq. (100 µM) MGO. These results show that there are only modest changes in glycation levels at R72 and R74 when using dialAGE$^{R42}$ mutations. As R72 is predicted to be somewhat proximal to Q49, these results suggest that the extent of dialAGE glycation modulation is sensitive to distance. Nondirectional (two-tailed) two-way ANOVA using Tukey's multiple comparison tests were used to determine if each variant yielded statistically significant differences in glycation. P<.05 (*), p<.01 (**), p<.001 (***), p<.0001 (****).

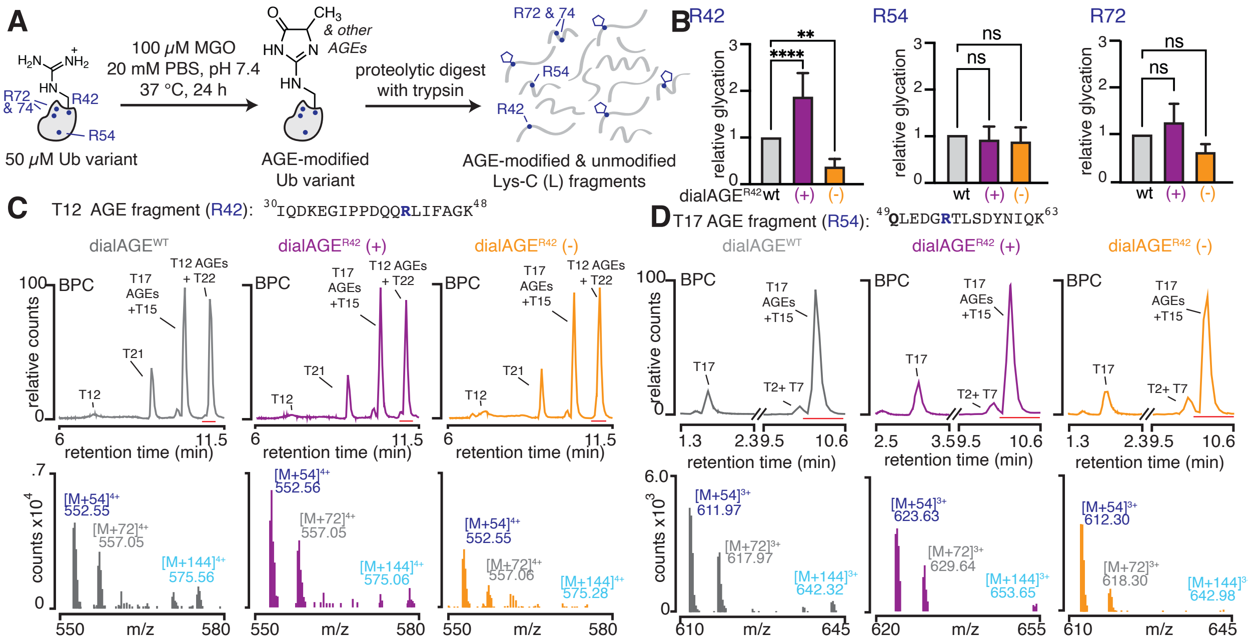

**Figure S7. DialAGE$^{R42}$ leads to site-specific differences in R42 glycation, as revealed with trypsin digest. A)** dialAGE$^{WT}$ and dialAGE$^{R42}$ variants (50 μM) were treated with 2 equiv. MGO (100 μM) and subsequently digested by trypsin. **B)** Quantification of relative glycation levels for each Arg reveal statistically significant differences solely at R42 for dialAGE$^{R42}$ Ub variants. Glycation was calculated using the same equation as with Lys-C, with peptides with up to 2 missed cleavages for both modified and unmodified fragments. **C)** Base peak chromatograms (BPC) (*top*) and combined spectra (*bottom*) over highlighted retention times showing increased glycation adducts for dialAGE$^{R42}$(+) and decreased glycation adducts for dialAGE$^{R42}$(-) at R42. **D)** BPC (*top*) and combined spectra at highlighted retention showing similar levels of modification for all variants at R54. As with Lys-C digests, these data show that mutations at Q49 produce site-specific differences in glycation at R42. Bar graphs represent mean ± SEM. Nondirectional (two-tailed) two-way ANOVA using Dunnett's multiple comparison tests were used to determine if each variant yielded statistically significant differences in glycation. P<.05 (*), p<.01 (**), p<.001 (***), p<.0001 (****).

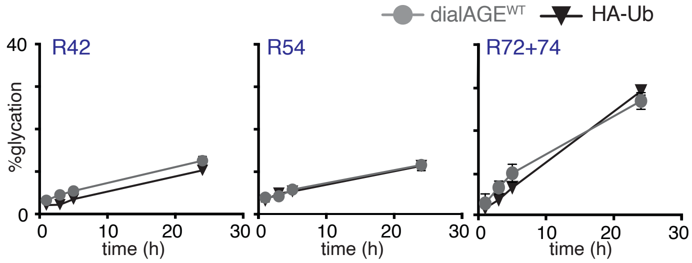


**Figure S8. DialAGE$^{WT}$ and commercially HA-Ub are similarly glycated.** Commercially available and dialAGE$^{WT}$ Ub (50 μM) were incubated with 2 eq. MGO over 24 h. Aliquots were taken at 1, 3, 5, and 24 h timepoints followed by Lys-C digestion. The % glycation at the resulting timepoints reveals no differences in glycation at each Arg between dialAGE$^{WT}$ Ub and commercially available HA-Ub constructs. Graph points represent mean ± SEM.

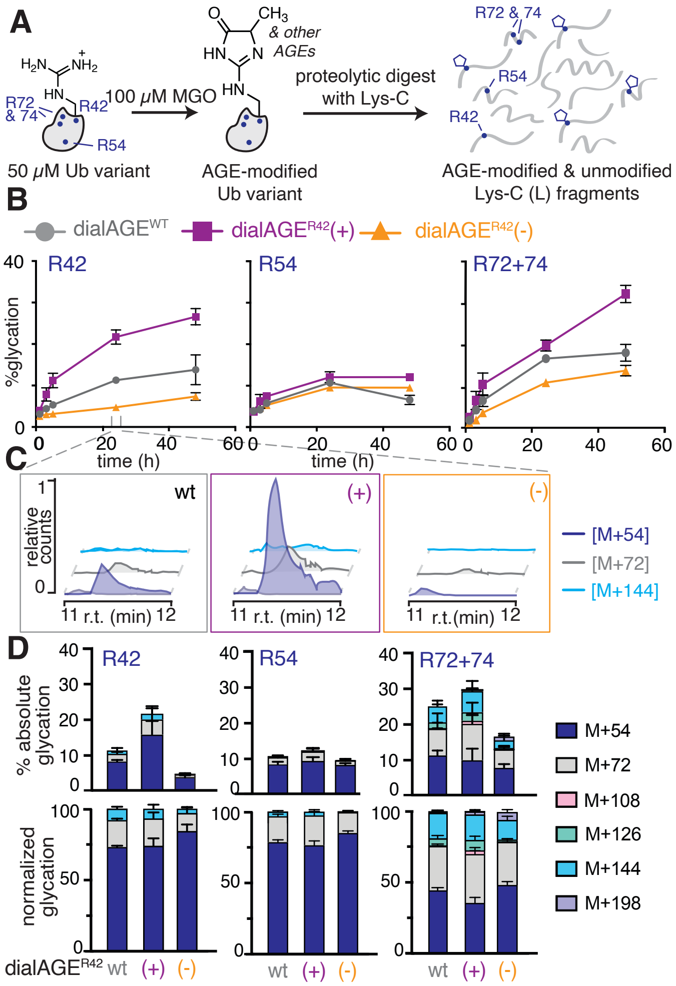


**Figure S9. DialAGE influences glycation rates.** **A)** To assess glycation levels as a function of time, dialAGE$^{WT}$ and dialAGE$^{R42}$ variants (50 μM) treated with 2 eq. MGO. Aliquots were removed at multiple timepoints (1, 3, 5, 24, 48 h), digested by Lys-C, and subsequently analyzed by LC-MS/MS **B)** Quantification of the % glycation for dialAGE$^{R42}$ variants over 48 h shows a consistent modulation of glycation at R42, even at early timepoints. Glycation at R54 remains unaltered across the three variants, while R72+74 exhibits small changes due to the proximity of R72 and Q49. **C)** Extracted ion chromatograms (EIC) reveal increased or decreased AGE formation, as shown for the 24 h time point. **D)** Quantification of the absolute and normalized AGE distributions show no major differences between AGEs formed for each dialAGE$^{R42}$ variant and dialAGE$^{WT}$, suggesting dialAGE works primarily by influencing rates of AGE formation. Bar graphs represent mean ± SEM.

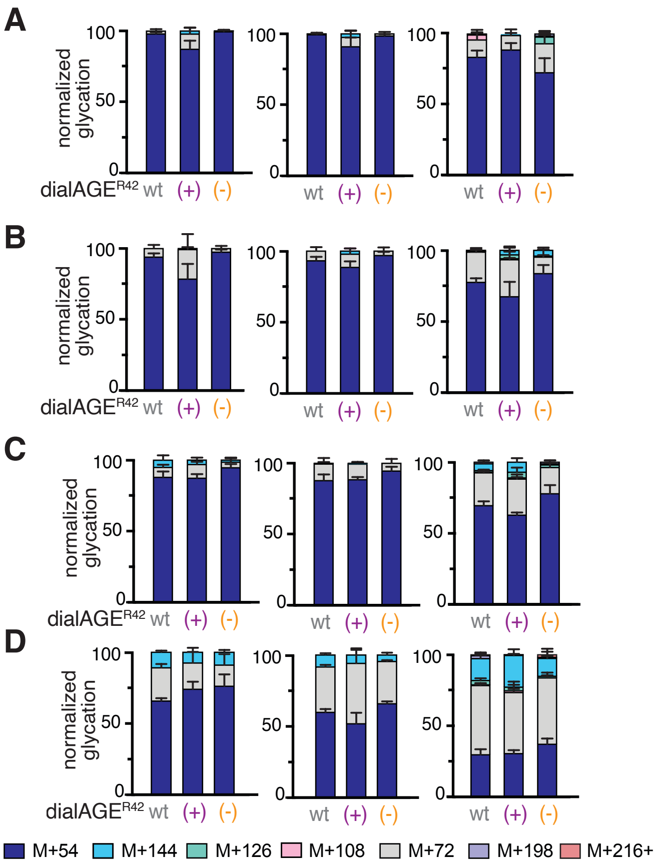


**Figure S10. DialAGE does not significantly modulate AGE distributions.** Normalized AGE distributions for dialAGE$^{R42}$ variants at (**A**) one, (**B**) three (**C**) five, and (**D**) 48 h show no major differences in the distribution of AGEs, suggesting that dialAGE$^{R42}$ alters only the rates of AGE formation and does not influence the preference for forming specific AGEs. Bar graphs represent mean ± SEM.

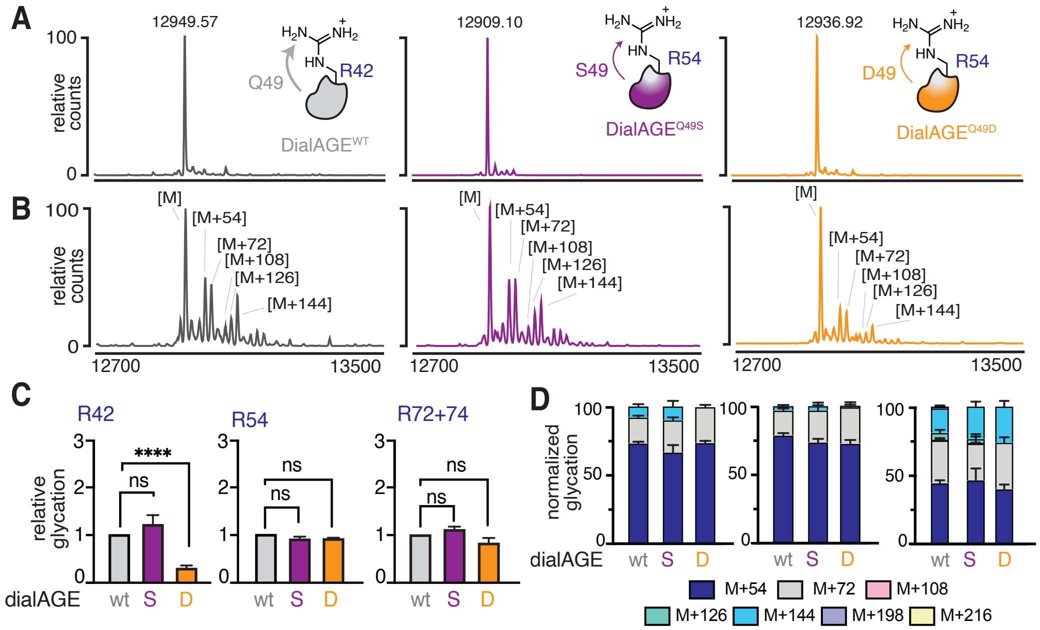


**Figure S11. Gln mutation to Asp leads to amplified decrease in R42 glycation.** Mass spectra of dialAGE$^{WT}$, dialAGE$^{R42}$(+)$^{Q49S}$, and dialAGE $^{R42}$(-)$^{Q49D}$ when incubated with **A)** 0 eq. MGO or **B)** 2 eq. MGO for 24 h. This results in no difference in modification for dialAGE$^{R42}$(+)$^{Q49S}$ compared to dialAGE$^{WT}$ and a decrease in modification for dialAGE$^{R42}$(-)$^{Q49D}$. **C)** Solely R42 glycation is hindered when dialAGE$^{R42}$(-)$^{Q49D}$ is compared to dialAGE$^{WT}$, however no Arg modification is modulated with dialAGE$^{R42}$(+)$^{Q49S}$ **D)** Normalized AGE distributions remain highly similar between all dialAGE$^{R42}$ variants, despite an absence of [M+144] adducts for dialAGE$^{R42}$(-)$^{Q49D}$. Bar graphs represent mean ± SEM. Nondirectional (two-tailed) two-way ANOVA using Dunnett's multiple comparison tests were used to determine if each variant yielded statistically significant differences in glycation. P<.05 (*), p<.01 (**), p<.001 (***), p<.0001 (****).

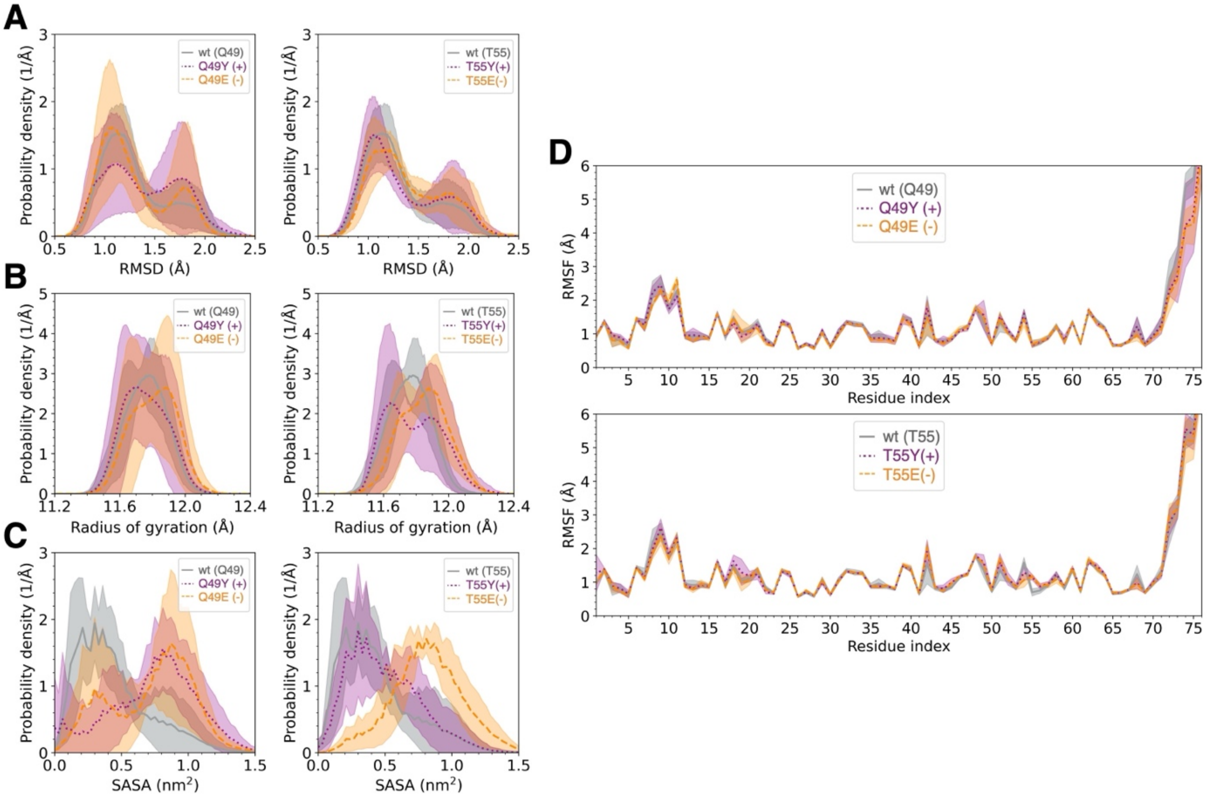


**Figure S12. Trends in root-mean-square deviation (RMSD), root-mean-square fluctuation (RMSF), radius of gyration, and solvent accessible surface area (SASA) cannot explain experimentally observed dialAGE behavior. A)** Average backbone RMSD for dialAGE$^{R42}$ (left) and dialAGE$^{R54}$ (right). Though dialAGE$^{R42}$ exhibits more variability compared to dialAGE$^{R54}$, no major difference is observed for dialAGE$^{R42}$ and dialAGE$^{R54}$ in comparison to dialAGE$^{WT}$. **B)** Average radius of gyration of protein for dialAGE$^{R42}$ (left) and dialAGE$^{R54}$ (right). Both dialAGE$^{R42}$(+) and dialAGE$^{R54}$(+) skew distributions to the left, and both dialAGE$^{R42}$(-) and dialAGE$^{R54}$(-) skew distributions to the right. The distributions of dialAGE$^{R54}$ are more spread in comparison to dialAGE$^{WT}$. These differences are minor in comparison to the generally overlapping distributions of all variants. **C)** Average solvent accessible surface area of R42 for dialAGE$^{R42}$ (left) and dialAGE$^{R54}$ (right). Both dialAGE$^{R42}$(+) and dialAGE$^{R42}$(-) increase SASA in comparison to dialAGE$^{WT}$. Meanwhile, dialAGE$^{R54}$(+) shows no major difference in comparison to dialAGE$^{WT}$, and dialAGE$^{R54}$(-) increases SASA in comparison to dialAGE$^{WT}$. **D)** Average RMSF per residue for dialAGE$^{R42}$ (left) and dialAGE$^{R54}$ (right). DialAGE$^{R42}$ shows no major difference in comparison to dialAGE$^{WT}$. Both dialAGE$^{R54}$(+) and dialAGE$^{R54}$(-) decrease fluctuations at R54 and increase fluctuations at site 55. All analyses are based on the last 100 ns trajectory of five parallel runs for each variant. The lines correspond to the average of the five parallel runs, and the shaded areas correspond to the standard deviations.

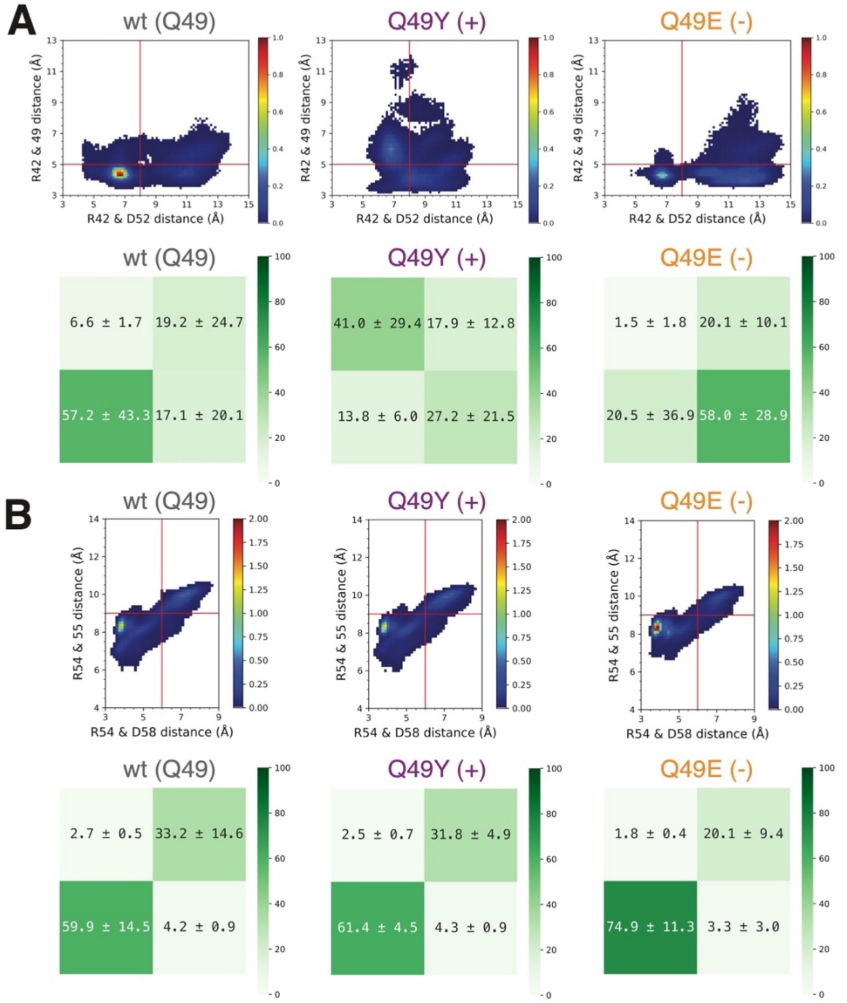


**Figure S13. DialAGE$^{R42}$ modulates distances surrounding R42 but not R54.**
**A)** Normalized 2D histograms (top) of R42…49 distances and R42…D52 distances show that dialAGE$^{R42}$(+) increased both R42…49 and R42…D52 distances, whereas dialAGE$^{R42}$(-) decreases R42…49 distances and increases R42…D52 distances. Then the histograms were divided into four quadrants, and the average and standard deviation of the population (bottom) for each quadrant were calculated. **B)** Normalized 2D histograms (top) of R54…55 distances and R54…D58 distances show that neither dialAGE$^{R42}$(+) nor dialAGE$^{R42}$(-) significantly modulates distances surrounding R54. The histograms were divided into four quadrants, and the average and standard deviation of the population (bottom) for each quadrant were calculated. The heatmap colors are based on the average values.

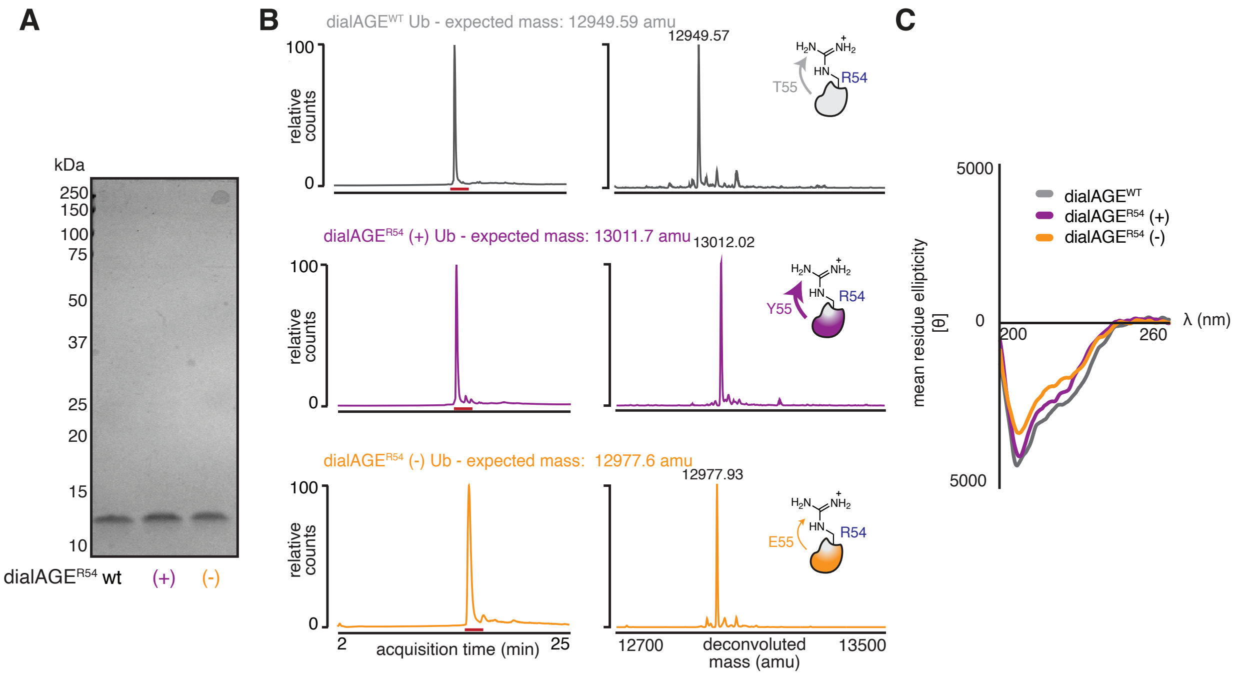


**Figure S14. Purification and characterization of dialAGE$^{R54}$ Ub variants**. Following expression and purification, dialAGE$^{R54}$ variants purity was assessed using (**A**) SDS-PAGE and (**B**) LC-MS, revealing >95% purity for all variants. **C**) Circular dichroism revealed no major differences in structure for dialAGE$^{R54}$ variants compared to dialAGE$^{WT}$.

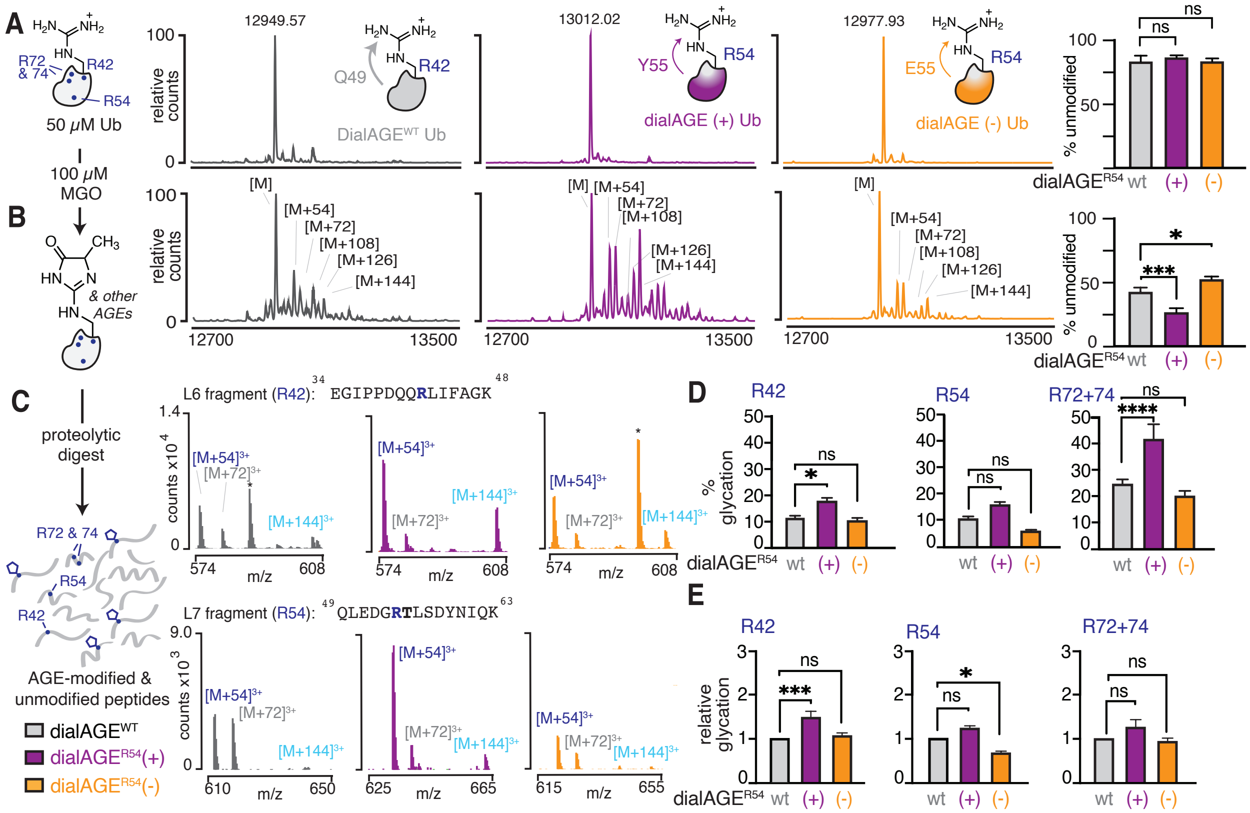


**Figure S15. Glycation modulation at R54 using dialAGE$^{R54}$. A**) Recombinant dialAGE$^{WT}$ and dialAGE$^{R54}$ variants (50 μM) were treated without or (**B**) with 2 equivalents of MGO (100 μM) for 24 h and analyzed by intact protein mass spectrometry. Total levels of glycation were determined by quantifying the intensity of the remaining unmodified peak ([M]) after MGO treatment. This resulted in increased or decreased levels of total protein glycation for dialAGE$^{R54}$(+) and (-) respectively. **C)** Combined spectra showing m/z for R42 (L6) and R54 (L7) modified fragments. These data show similar levels of R42 glycation, but decreased R54 glycation for dialAGE$^{R54}$(-). For dialAGE$^{R54}$(+) increased levels of glycation were observed at both sites. * denotes a variant-specific fragment that coelutes with the L6 fragment only for dialAGE$^{R54}$(-). **D**) Quantification of the absolute % glycation for dialAGE$^{R54}$ variants at each Arg. **E**) Relative glycation at each Arg for each dialAGE$^{R54}$ variant. These results reveal an unexpected global increase in glycation at all Arg for dialAGE$^{R54}$(+), but the expected decrease in glycation solely at R54 for dialAGE$^{R54}$(-). Bar graphs represent mean ± SEM. Nondirectional (two-tailed) two-way ANOVA using Tukey's multiple comparison tests were used to determine if each variant yielded statistically significant differences in glycation. P<.05 (*), p<.01 (**), p<.001 (***), p<.0001 (****).

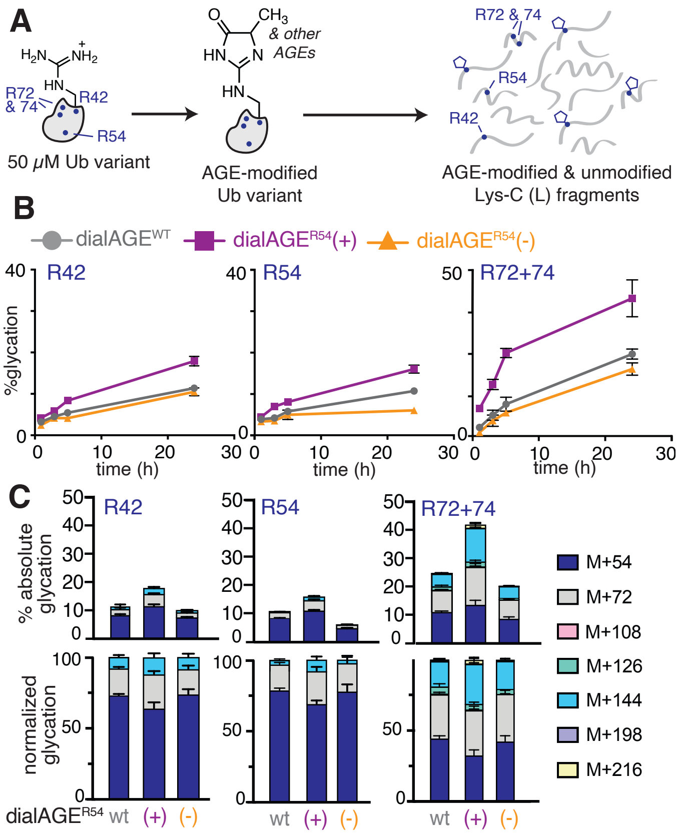


**Figure. S16. DialAGE$^{R54}$ modulates glycation levels but not AGE distributions. A)** dialAGE$^{WT}$ and dialAGE$^{R54}$ variants (50 µM) were treated with 2 equiv. MGO (100 µM). Aliquots were removed at multiple timepoints (1, 3, 5, 24, 48 h), digested by Lys-C, and subsequently analyzed by LC-MS/MS **B)** Quantification of the % glycation for dialAGE$^{R54}$ variants over 48 h shows that dialAGE$^{R54}$(-) decreases R54 glycation, while dialAGE$^{R54}$(+) increases it. Glycation of R42 and R72+74 was unaffected by dialAGE$^{R54}$(-), while both consistently exhibit increased glycation for dialAGE$^{R54}$(+), even at the earliest timepoints. **C)** Comparison of absolute glycation (*top*) and relative AGE distributions (*bottom*) after 24 h of MGO incubation show no major differences between AGE distributions of dialAGE$^{R54}$ variants. Graphs represent mean ± SEM.

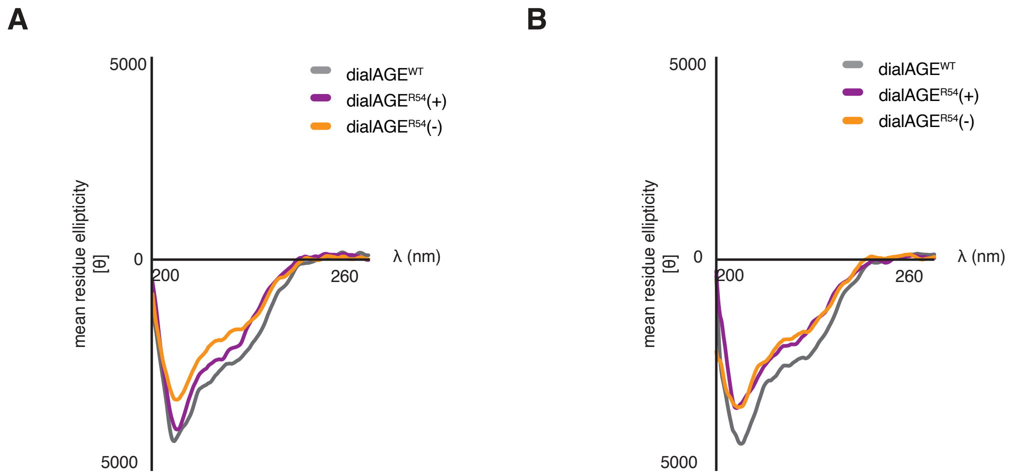


**Figure S17. Circular dichroism at elevated temperature reveals no structural differences in dialAGE$^{R54}$ variants.** Representative circular dichroism spectra obtained at (**A**) the temperature at which glycation reactions are performed and (**B**) an elevated 60 ºC. These spectra show no significant differences, suggesting that the elevated levels of glycation observed for dialAGE$^{R54}$(+) is not due to decreased thermal stability and structural unfolding.

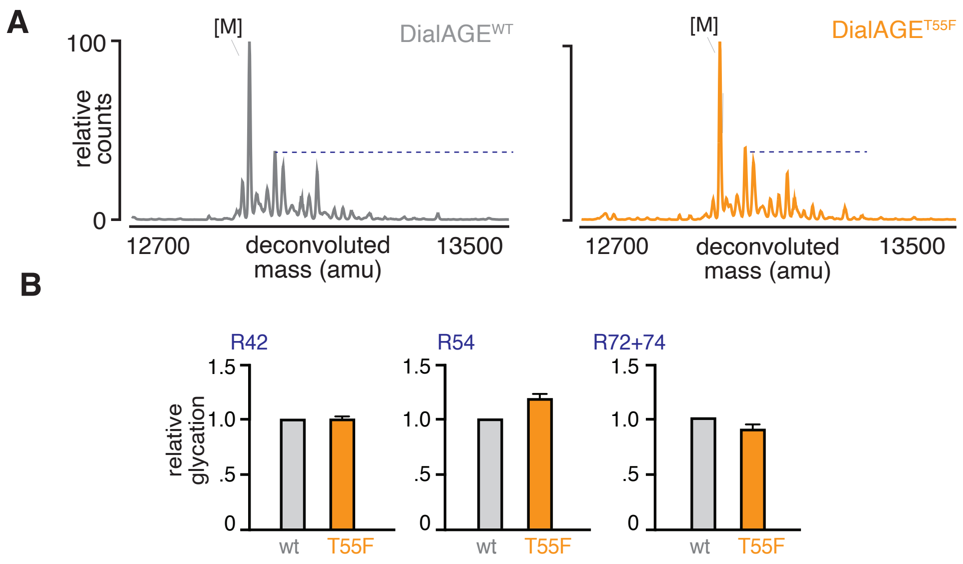


**Figure S18. Phenylalanine mutation at T55 does not impact glycation. A**) Representative intact mass spectra display no differences in total protein glycation between dialAGE$^{WT}$ and dialAGE$^{R54}$(-)$^{T55F}$. **B**) Relative glycation for each Arg fragment reveals no differences in glycation at any Arg. Bar graphs represent mean ± SEM.

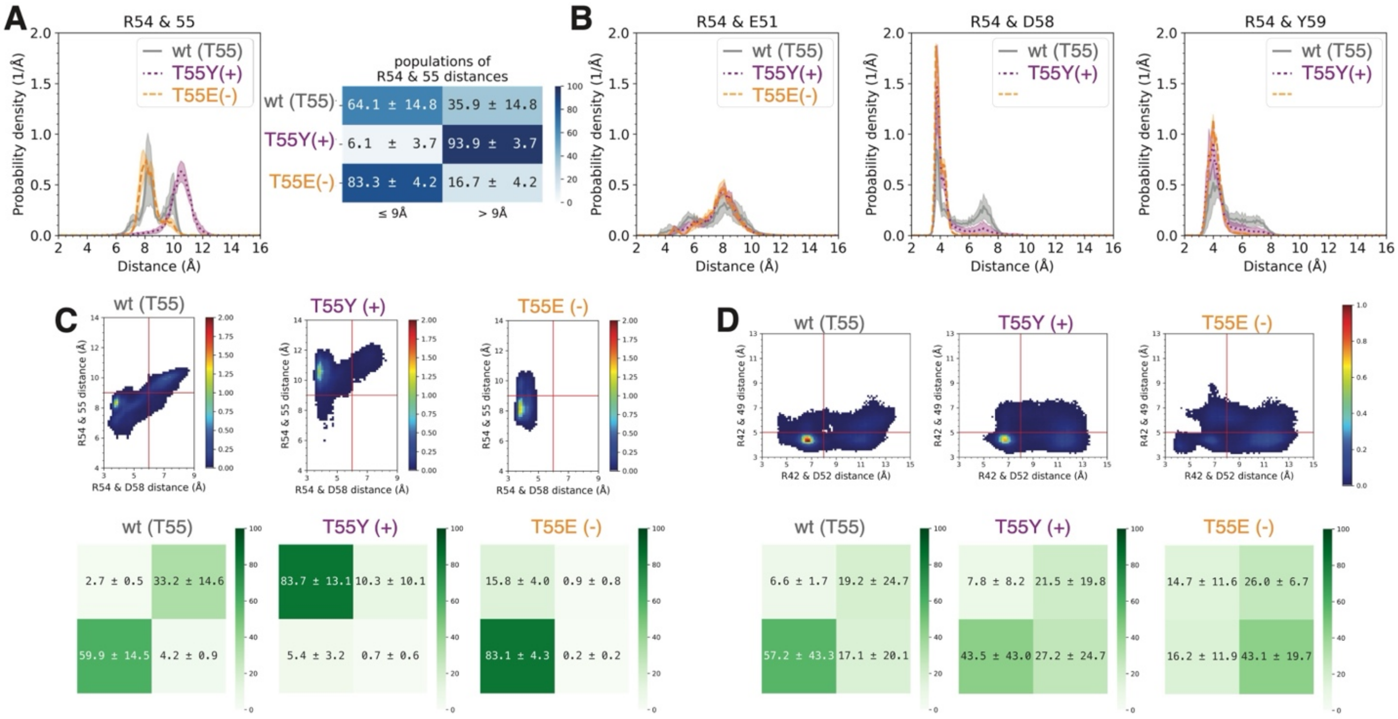


**Figure S19. Molecular dynamics simulations for dialAGE$^{R54}$. A)** Distance distributions (with standard deviations in shaded regions) between the terminal carbon of the R54 side chain and the terminal carbon of the side chain at site 55 (left). The average and standard deviation of the population with R54…55 distances ≤ 9Å and >9 Å were calculated (heatmap colors correspond to the average values). **B)** Distance distributions (with standard deviations in shaded regions) between the terminal carbon of R54 side chain and the terminal carbon of the E51 (left), D58 (middle), and Y59 (right). **C)** Normalized 2D histograms (top) of R54…55 distances and R54…D58 distances show that dialAGE$^{R54}$(+) increases R54…55 distances and dialAGE$^{R54}$(+) decreases R54…55 distances. The histograms were then divided into four quadrants, and the average and standard deviation of the population (bottom) for each quadrant were calculated. **D)** Normalized 2D histograms (top) of R42…49 distances and R42…D52 distances show that dialAGE$^{R54}$(+) does not significantly modulate R42…49 distances but slightly increases R42…D52 distances. DialAGE$^{R54}$(+) increases both R42…49 distances and R42…D52 distances. The histograms were divided into four quadrants, and the average and standard deviation of the population (bottom) for each quadrant were calculated. The heatmap colors are based on the average values.

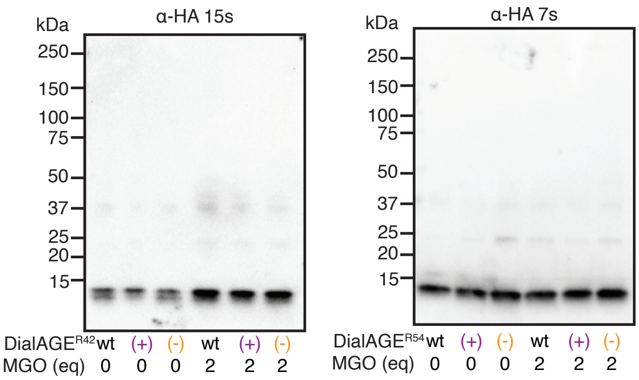


**Figure S20. MGO treatment does not change protein concentrations.** Following treatment of dialAGE$^{R42}$ (left) and dialAGE$^{R54}$ (right) variants with 2 eq. MGO for 24 h, the reactions were quenched and aliquots analyzed by western blotting using α-HA primary antibody. The protein signal in the resulting blots was consistent between treated and untreated samples for all variants.

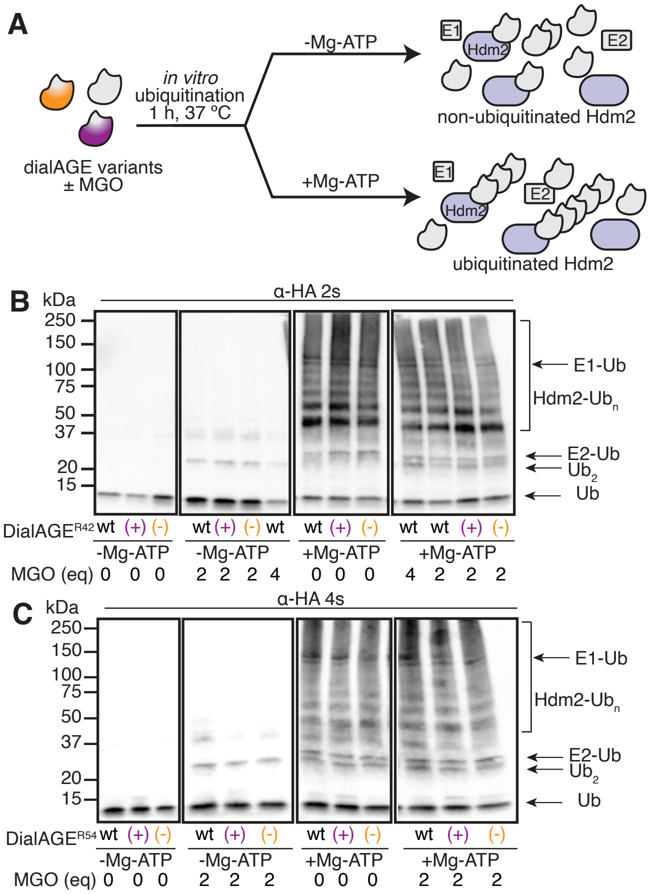


**Figure S21. *In vitro* ubiquitination using dialAGE Ub variants A**) To evaluate the impact of dialAGE on Ub function, an *in vitro* ubiquitination assay was used. DialAGE variants were treated ± MGO for 24 h, and subsequently incubated with E1 (His6-Ube1), E2 (UbE2D1), and E3 (GST-Hdm2 RING domain) enzymes, either with or without Mg2+-ATP for 1 h. The resulting polyubiquitinated proteins were analyzed by western blot, tracking the HA tag on (**B**) dialAGE$^{R42}$ or (**C**) and dialAGE$^{R54}$ variants. These data show uniform incorporation of unmodified or glycated dialAGE variants into Ub ladders, suggesting glycation does not impact the ability of ubiquitin to be conjugated onto UPS substrates.

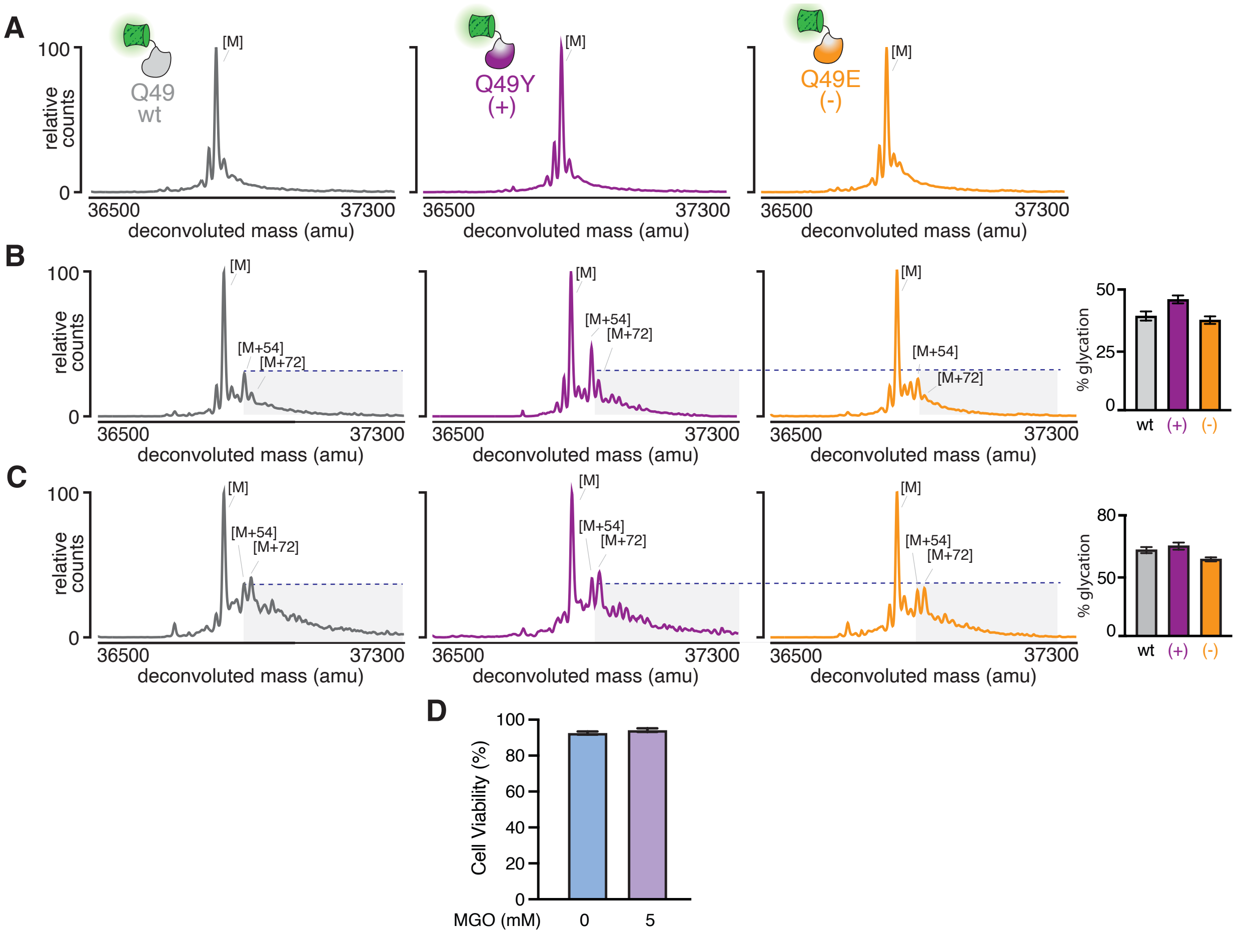


**Figure S22. Ub-GFP dialAGE$^{R42}$ variants have scaled levels of glycation with increasing MGO**. Intact mass spectra of dialAGE$^{R42}$ Ub-GFP variants following a 2 hr treatment with (**A**) 0 mM MGO, (**B**) 5 mM MGO, and (**C**) 7 mM MGO. Following each of the MGO treatment conditions there are observable differences in glycation between dialAGE$^{R42}$ variants by comparison of [M+54] peaks compared to the unmodified [M] peak. Treatment with 7 mM MGO increases levels of modification modestly compared to the 5 mM MGO treatment condition, however, differences in levels of glycation are decreased. **D)** Cell viability as measured by a trypan blue assay results in highly consistent cell viability between cells treated with 0 and 5 mM MGO for 2 h.

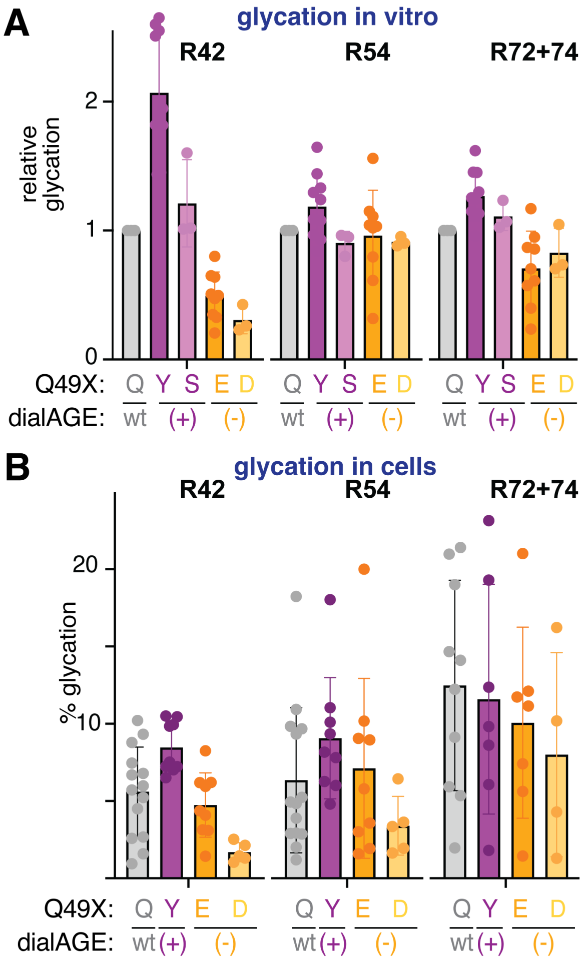


**Figure S23. Ub-GFP dialAGE$^{R42}$(-)$^{Q49D}$ leads to greater dynamic range in glycation signal**. **A)** Comparison of relative levels of glycation observed *in vitro* for dialAGE$^{WT}$ and the four dialAGE$^{R42}$ variants reveals that while only dialAGE$^{R42}$(+) leads to marked increased glycation at R42, dialAGE$^{R42}$(-)$^{Q49D}$ hinders glycation further at R42 compared to dialAGE$^{R42}$(-), suggesting Asp may be a better acidic residue to prevent glycation than Glu. However, none of the four dialAGE$^{R42}$ variants produced differences in glycation at R54, and only minimal difference at R72+74. **B)** Following treatment of cells with 5 mM MGO, Ub-GFP dialAGE$^{R42}$(-)$^{Q49D}$ displays a further decreased level of glycation at R42 compared to Ub-GFP dialAGE$^{WT}$. Although glycation signal at the R54 and R72+74 sites was more variable in general, they are still comparable to that observed for Ub-GFP dialAGE$^{WT}$. Together, these data suggest that the use of Q49D, rather than Q49E for dialAGE(-) is likely to provide a greater dynamic range for glycation at R42, enabling the study of the effect of R42 glycation in ubiquitin biology.

**Table S1. Ub DialAGE Lys-C fragments**
Lys-C fragment annotation used in labeled BPC peaks for Lys-C digests.

Lys-C Fragments

| Fragment | Missed Cleavages | Reactive Arg | Sequence |
|---|---|---|---|
| L1 | 0 | | SYYHHHHHHLESTSLYK |
| L2 | 0 | | AGFTMGSMYPYDVPDYASGSMQIFVK |
| L3 | 0 | | TLTGK |
| L4 | 0 | | TITLEVEPSDTIENVK |
| L5 | 0 | | IQDK |
| L6 | 0 | R42 | EGIPPDQQ**R**LIFAGK |
| L7 | 0 | R54 | QLEDG**R**TLSDYNIQK |
| L8 | 0 | R72+R74 | ESTLHLVL**R**L**R**GG |
| L9 | 1 | | SYYHHHHHHLESTSLYKK |
| L10 | 1 | | KAGFTMGSMYPYDVPDYASGSMQIFVK |
| L11 | 1 | | AGFTMGSMYPYDVPDYASGSMQIFVKTLTGK |
| L12 | 1 | | TLTGKTITLEVEPSDTIENVK |
| L13 | 1 | | TITLEVEPSDTIENVKAK |
| L14 | 1 | | AKIQDK |
| L15 | 1 | R42 | IQDKEGIPPDQQ**R**LIFAGK |
| L16 | 1 | R42+R54 | EGIPPDQQ**R**LIFAGKQLEDG**R**TLSDYNIQK |
| L17 | 1 | R72+R74 | QLEDGRTLSDYNIQKESTLHLVL**R**L**R**GG |
| L18 | 2 | | SYYHHHHHHLESTSLYKKAGFTMGSMYPYDVPDYASGSMQIFVK |
| L19 | 2 | | KAGFTMGSMYPYDVPDYASGSMQIFVKTLTGK |
| L20 | 2 | | AGFTMGSMYPYDVPDYASGSMQIFVKTLTGKTITLEVEPSDTIENVK |
| L21 | 2 | | TLTGKTITLEVEPSDTIENVKAK |
| L22 | 2 | | TITLEVEPSDTIENVKAKIQDK |
| L23 | 2 | R42 | AKIQDKEGIPPDQQ**R**LIFAGK |
| L24 | 2 | R42+R54 | IQDKEGIPPDQQ**R**LIFAGKQLEDG**R**TLSDYNIQK |
| L25 | 2 | R42, R54, R72, R74 | EGIPPDQQ**R**LIFAGKQLEDG**R**TLSDYNIQKESTLHLVL**R**L**R**GG |

**Table S2. Ub DialAGE tryptic fragments**
Tryptic fragment annotation used in labeled BPC peaks for tryptic digests.

Tryptic Fragments

| Fragment | Missed Cleavages | Reactive Arg | Sequence |
|---|---|---|---|
| T1 | 0 | | SYYHHHHHHLESTSLYK |
| T2 | 1 | | SYYHHHHHHLESTSLYKK |
| T3 | 1 | | KAGFTMGSMYPYDVPDYASGSMQIFVK |
| T4 | 0 | | AGFTMGSMYPYDVPDYASGSMQIFVK |
| T5 | 1 | | AGFTMGSMYPYDVPDYASGSMQIFVKTLTGK |
| T6 | 0 | | TLTGK |
| T7 | 1 | | TLTGKTITLEVEPSDTIENVK |
| T8 | 0 | | TITLEVEPSDTIENVK |
| T9 | 1 | | TITLEVEPSDTIENVKAK |
| T10 | 1 | | AKIQDK |
| T11 | 0 | | IQDK |
| T12 | 1 | R42 | IQDKEGIPPDQQ**R** |
| T13 | 0 | R42 | EGIPPDQQ**R** |
| T14 | 1 | R42 | EGIPPDQQ**R**LIFAGK |
| T15 | 0 | | LIFAGK |
| T16 | 1 | R54 | LIFAGKQLEDG**R** |
| T17 | 0 | R54 | QLEDG**R** |
| T18 | 1 | R54 | QLEDG**R**TLSDYNIQK |
| T19 | 0 | | TLSDYNIQK |
| T20 | 1 | R72 | TLSDYNIQKESTLHLVL**R** |
| T21 | 0 | R72 | ESTLHLVL**R** |
| T22 | 1 | R72 + R74 | ESTLHLVL**R**L**R** |
| T23 | 0 | R74 | L**R** |
| T24 | 1 | R74 | L**R**GG |

### Table S3. Expected and observed fragments for R42

| R42 | L6 fragment | | DialAGE$^{WT}$ | | |
|---|---|---|---|---|---|
| | Fragment MW | Expected m/z | Observed fragment MW | observed m/z | error (ppm) |
| [M] | 1667.8995 | 556.9665 | 1667.9009 | 556.9669667 | -0.837872056 |
| [M+54] | 1721.9095 | 574.9698333 | 1721.9054 | 574.9684667 | 2.376936297 |
| [M+72 | 1739.9195 | 580.9731667 | 1739.9216 | 580.9738667 | -1.204874924 |
| [M+144] | 1811.9395 | 604.9798333 | 1811.9348 | 604.9782667 | 2.589618001 |

| | L6 fragment | | dialAGE$^{R42}$(+) | | |
|---|---|---|---|---|---|
| | Fragment MW | Expected m/z | Observed fragment MW | observed m/z | error (ppm) |
| [M] | 1667.8995 | 556.9665 | 1667.8987 | 556.9662333 | 0.478784032 |
| [M+54] | 1721.9095 | 574.9698333 | 1721.9083 | 574.9694333 | 0.695688672 |
| [M+72 | 1739.9195 | 580.9731667 | 1739.9174 | 580.9724667 | 1.204874924 |
| [M+144] | 1811.9395 | 604.9798333 | 1811.9375 | 604.9791667 | 1.101965107 |

| | L6 fragment | | dialAGE$^{R42}$(-) | | |
|---|---|---|---|---|---|
| | Fragment MW | Expected m/z | Observed fragment MW | observed m/z | error (ppm) |
| [M] | 1667.8995 | 556.9665 | 1667.8964 | 556.9654667 | 1.855288125 |
| [M+54] | 1721.9095 | 574.9698333 | 1721.9062 | 574.9687333 | 1.913143849 |
| [M+72 | 1739.9195 | 580.9731667 | 1739.9195 | 580.9731667 | 0 |
| [M+144] | 1811.9395 | 604.9798333 | 1811.9278 | 604.9759333 | 6.446495875 |

### Table S4. Expected and observed fragments for R54

| **R54** | **L7 fragment** | | DialAGE$^{WT}$ | | |
|---|---|---|---|---|---|
| | Fragment MW | Expected m/z | Observed fragment MW | observed m/z | error (ppm) |
| [M] | 1778.8799 | 593.9599667 | 1778.883 | 593.961 | -1.7397357 |
| [M+54] | 1832.8899 | 611.9633 | 1832.8893 | 611.9631 | 0.32681698 |
| [M+72 | 1850.8999 | 617.9666333 | 1850.9012 | 617.9670667 | -0.7012245 |
| [M+144] | 1922.9199 | 641.9733 | 1922.9112 | 641.9704 | 4.51732183 |

| | **L7 fragment** | | dialAGE$^{R42}$(+) | | |
|---|---|---|---|---|---|
| | Fragment MW | Expected m/z | Observed fragment MW | observed m/z | error (ppm) |
| [M] | 1813.8846 | 605.6282 | 1813.8863 | 605.6287667 | -0.9356676 |
| [M+54] | 1867.8946 | 623.6315333 | 1867.8922 | 623.6307333 | 1.28280877 |
| [M+72 | 1885.9046 | 629.6348667 | 1885.9087 | 629.6362333 | -2.1705702 |
| [M+144] | 1957.9246 | 653.6415333 | 1957.9237 | 653.6412333 | 0.45896716 |

| | **L7 fragment** | | dialAGE$^{R42}$(-) | | |
|---|---|---|---|---|---|
| | Fragment MW | Expected m/z | Observed fragment MW | observed m/z | error (ppm) |
| [M] | 1779.864 | 594.288 | 1779.857 | 594.2856667 | 3.92626695 |
| [M+54] | 1833.874 | 612.2913333 | 1833.8698 | 612.2899333 | 2.28649325 |
| [M+72 | 1851.884 | 618.2946667 | 1851.8799 | 618.2933 | 2.21038081 |
| [M+144] | 1923.904 | 642.3013333 | 1923.8998 | 642.2999333 | 2.1796623 |

### Table S5. Expected and observed fragments for R72+74

| R72+74 | L8 fragment | | dialAGE$^{WT}$ | | |
|---|---|---|---|---|---|
| | Fragment MW | Expected m/z | Observed fragment MW | observed m/z | error (ppm) |
| [M] | 1449.8416 | 484.2805333 | 1449.8446 | 484.2815333 | -2.0649188 |
| [M+54] | 1503.8516 | 502.2838667 | 1503.8492 | 502.2830667 | 1.59272486 |
| [M+72 | 1521.8616 | 508.2872 | 1521.8638 | 508.2879333 | -1.4427539 |
| [M+108] | 1557.8612 | 520.2870667 | 1557.8644 | 520.2881333 | -2.0501503 |
| [M+126] | 1575.8708 | 526.2902667 | 1575.8726 | 526.2908667 | -1.1400553 |
| [M+144] | 1593.8816 | 532.2938667 | 1593.8879 | 532.2959667 | -3.9451892 |

| R72+74 | L8 fragment | | dialAGE$^{R42}$(+) | | |
|---|---|---|---|---|---|
| | Fragment MW | Expected m/z | Observed fragment MW | observed m/z | error (ppm) |
| [M] | 1449.8416 | 484.2805333 | 1449.8424 | 484.2808 | -0.550645 |
| [M+54] | 1503.8516 | 502.2838667 | 1503.8524 | 502.2841333 | -0.5309083 |
| [M+72 | 1521.8616 | 508.2872 | 1521.8618 | 508.2872667 | -0.1311594 |
| [M+108] | 1557.8612 | 520.2870667 | 1557.8671 | 520.2890333 | -3.7799645 |
| [M+126] | 1575.8708 | 526.2902667 | 1575.8651 | 526.2883667 | 3.61017507 |
| [M+144] | 1593.8816 | 532.2938667 | 1593.8838 | 532.2946 | -1.3776851 |

| R72+74 | L8 fragment | | dialAGE$^{R42}$(-) | | |
|---|---|---|---|---|---|
| | Fragment MW | Expected m/z | Observed fragment MW | observed m/z | error (ppm) |
| [M] | 1449.8416 | 484.2805333 | 1449.84 | 484.28 | 1.10129005 |
| [M+54] | 1503.8516 | 502.2838667 | 1503.8517 | 502.2839 | -0.0663635 |
| [M+72 | 1521.8616 | 508.2872 | 1521.8595 | 508.2865 | 1.37717416 |
| [M+108] | 1557.8612 | 520.2870667 | 1557.845 | 520.2816667 | 10.3788857 |
| [M+126] | 1575.8708 | 526.2902667 | 1575.864 | 526.288 | 4.30687552 |
| [M+144] | 1593.8816 | 532.2938667 | 1593.8825 | 532.2941667 | -0.5635985 |
